# A framework for genomics-informed ecophysiological modeling in plants

**DOI:** 10.1101/497537

**Authors:** Diane R. Wang, Carmela R. Guadagno, Xiaowei Mao, D. Scott Mackay, Jonathan R. Pleban, Robert L. Baker, Cynthia Weinig, Jean-Luc Jannink, Brent E. Ewers

**Author notes:** Corresponding author:Diane R. Wang. Co-first authors (Diane R. Wang, Carmela R. Guadagno, Xiaowei Mao).

## Abstract

Dynamic process-based plant models capture complex physiological response across time, carrying the potential to extend simulations out to novel environments and lend mechanistic insight to observed phenotypes. Despite the translational opportunities for varietal crop improvement that could be unlocked by linking natural genetic variation to first-principles based modeling, these models are challenging to apply to large populations of related individuals. Here we use a combination of model development, experimental evaluation, and genomic prediction in *Brassica rapa* L. to set the stage for future large-scale process-based modeling of intra-specific variation. We develop a new canopy growth sub-model for *B. rapa* within the process-based model Terrestrial Regional Ecosystem Exchange Simulator (TREES), test input parameters for feasibility of direct estimation with observed phenotypes across cultivated morphotypes and indirect estimation using genomic prediction on a Recombinant Inbred Line population, and explore model performance on an *in silico* population under non-stressed and mild water stressed conditions. We find evidence that the updated whole plant model has capacity to distill genotype by environment interaction (G x E) into tractable components. The framework presented offers a means to link genetic variation with environment-modulated plant response and serves as a stepping stone towards large-scale prediction of un-phenotyped, genetically-related individuals under un-tested environmental scenarios.

## INTRODUCTION

Beginning in the 1960s, dynamic process-based models for plants have been implemented successfully to address research questions and inform decision-making across a variety of fundamental and applied settings (Alberda and Sibma, 1968; de Wit et al., 1970; Hoogenboom, 2000; Jones et al., 2003; Keating et al., 2003; Sinclair et al., 2010; Messina et al., 2015; Thorp et al., 2017; Johnson et al., 2018). These models enable studying time-series responses of a plant system to various inputs by formalizing physiological processes into assemblies of mathematical functions. While a diversity of tools exist, with names such as ‘biophysical models’, ‘ecophysiological models’, and ‘crop growth models’, one unifying feature is that they can readily synthesize frontier knowledge from divergent disciplines (Bouman et al., 1996), thereby adapting continuously to a changing research landscape. Since the rise of the genomics era, the capacity of process-based modeling to handle temporal climatic variation and extend predictions to novel scenarios such as stress conditions has made it attractive to scientific communities interested in dissecting the genetics of environmentally-plastic traits or in predicting performance for breeding (White and Hoogenboom, 1996; White and Hoogenboom, 2003; Hammer et al., 2005; Hammer et al., 2006; Messina et al., 2006; Struik, 2012; Yin et al., 2018). Despite the translational opportunities proffered by successful linking of genetic variation and physiological dynamics via process-based modeling, major challenges for extending these models to handle the sizable populations relevant to plant geneticists and breeders remain. How can we systematically generate large sets of genotype-specific input parameters needed by biophysical models? Literature values available at species or genus level often used for modeling may be suitable for conserved traits with little intraspecific variation but are not appropriate for traits expressing high variability at the genotype-level. Conversely, how does process model structure need to be modified to utilize genotype-specific inputs? First-principles based models are conventionally developed without considerable evaluation for the genetic relevance of parameters.

Innovative approaches to address the challenge of combining process modeling and natural genetic variation have been investigated with varying degrees of success. Early efforts involved modeling the effects of quantitative trait loci (QTL) discovered in mapping populations to inform process model parameters or using modeling outcomes for QTL discovery itself (Yin et al., 1999; Yin, 2000; Yin et al., 2005). Another early example is the incorporation of seven genes that affected phenology, growth habit, and seed size into BEANGRO, a process model for the common bean, *Phaseolus vulgaris* L (White and Hoogenboom, 1996). One downside to a gene or QTL-based approach for process model parameterization is that typically only loci with large or moderate effect sizes can be used as predictors because loci of small effects are likely to be missed in the discovery process. An alternative is genomic prediction (GP), which entails estimating breeding values or other traits of un-phenotyped individuals using whole-genome models trained on related phenotyped individuals (Meuwissen et al., 2001). Compared with QTL-based approach, genetic effects are estimated for each marker regardless of effect sizes, and then are summed up to predict the overall breeding value. GP was first implemented in cattle breeding to increase rate of genetic gain (Schaeffer, 2006) and has since been adopted across plant breeding communities for black-box (empirical) prediction of agronomically important traits (Heffner et al., 2011; Rutkoski et al., 2015; Spindel et al., 2015; Thavamanikumar et al., 2015; Minamikawa et al., 2017; Okeke et al., 2017; Wolfe et al., 2017). Recently, Technow et al. developed a method to capture biological information in predictions by integrating crop modeling directly into a genomic prediction framework, which was later applied to a maize dataset (Technow et al., 2015; Cooper et al., 2016). The reverse application, i.e. utilization of genomic prediction to inform model parameters has been proposed (Tardieu et al., 2018) but to our knowledge it has not yet been implemented.

Here we explore a modeling framework in *Brassica rapa* L. that leverages genomic prediction to drive genotype-dependent variability of a plant process model, Terrestrial Regional Ecosystem Exchange Simulator (TREES) (Mackay et al., 2015). This model was selected for its capacity to model rhizosphere and xylem hydraulics, critical processes in drought stressed conditions. Recognizing that growth and development are likely to be highly genotype-specific, we build and iteratively update a new canopy growth sub-model within TREES as the principal link between genetic complexity and dynamic physiological response, testing it under non-stressed and water-limited scenarios. We ask several fundamental questions: Can input parameters be directly estimated from real-world observation? How does uncertainty at the parameter level propagate to outcome uncertainty? Are parameters genetically relevant and can they then be predicted using genomic information?

## MATERIALS AND METHODS

### Model development

The process model TREES couples soil-plant hydraulics with plant physiological routines such as photosynthesis and carbon allocation. The model has been well-tested under various natural and experimental conditions primarily for long-lived woody species (Mackay et al., 2015; McDowell et al., 2016; Tai et al., 2017; Johnson et al., 2018), where development was assumed to occur across very long timeframes (e.g., years or decades). In order to extend the application of TREES for use in genotypes of short-lived herbaceous plants that express dramatic ontogenetic changes over relatively short timeframes (e.g., days, weeks, and months), we developed a new sub-model to capture temperature-dependent canopy growth based on the emergence and expansion events of individual leaves. The other sub-processes within TREES were retained, with the rationale that the mechanisms underlying fundamental biophysical processes such as C3 photosynthesis and hydraulics are conserved and therefore can be dealt with via proper parameterization versus a design overhaul. The new canopy sub-model uses 22 additional input parameters (**Table 1**), selected based on feasibility of being either empirically measured or predicted with genomic models, and is designed to begin model simulations at the initiation of the third true leaf. The primary governing equation used for computation of theoretical relative rate of leaf growth:

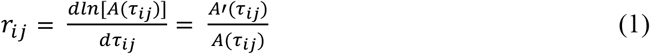

where *τ_ij_* is thermal time of leaf *i* spent in process *j* (cell division or leaf expansion). *A*(*τ_ij_*) is the thermal time-dependent leaf area following a logistic growth function

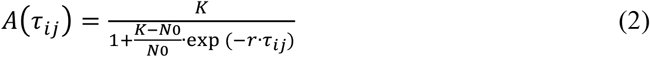

parameterized by three inputs *K, N_0_*, and *r*. Details on model development including other governing equations are presented in **Appendix S1** and a pipeline describing how to derive leaf area parameter information for the new sub-model in TREES is presented in **Figure S1**. Scripts associated with the pipeline may be accessed at https://github.com/DRWang3/leaf_model_TREES_paper along with the version of TREES used in this study. For sensitivity analysis simulations of stochastic leaf growth parameters, we sampled from distributions for all three leaf parameters, *K, N_0_,* and *r*, at either low or high uncertainty levels (40^th^-60^th^ quantile and 10-90^th^ quantile, respectively). Original TREES parameters were held invariant across the genotypes.

**Table 1.** Input parameters for the new canopy growth sub-model

| Type | Parameter | Description | Unit |
| --- | --- | --- | --- |
| module on/off | useLeafModule | 1 to grow individual leaves; 0 otherwise | unitless |
| leaf growth and development | leafAreaMax* | Parameter K from logistic growth function | cm<sup>2</sup> |
|  | initialLeafSize* | Parameter N0 from logistic growth function | cm<sup>2</sup> |
|  | leafArea_Rate* | Parameter r from logistic growth function | unitless |
|  | useLeafGamma | 1 for stochastic variation in leaf growth; 0 for deterministic simulation | unitless |
|  | Kalpha* | shape parameter of gamma distribution for leafAreaMax | unitless |
|  | Kbeta* | rate parameter of gamma distribution for leafAreaMax | unitless |
|  | Nalpha* | shape parameter of gamma distribution for initialLeafSize | unitless |
|  | Nbeta* | rate parameter of gamma distribution for initialLeafSize | unitless |
|  | ralpha* | shape parameter of gamma distribution for leafArea_Rate | unitless |
|  | rbeta* | rate parameter of gamma distribution for leafArea_Rate | unitless |
|  | dur_LeafExpansion | thermal time to 95% of leafAreaMax | °C hr |
|  | SLA_low | genotype-specific lower limit for specific leaf area | kg C/m<sup>2</sup> |
|  | leaf_insertAngle | angle between leaf petiole and main stem | degree |
|  | leaf_len_to_width | ratio between leaf length and width | unitless |
| temperature-dependent development | phyllochron | thermal time between leaf initiation events | °C hr |
|  | floweringTime | thermal time to flowering | °C hr |
|  | Tbase | developmental base temperature | °C |
| model initiation | therm_plant_init | thermal time to third leaf initiation (approximated by leaf emergence) | °C hr |
|  | projectedArea_init | projected shoot area at third leaf initiation (approximated by leaf emergence) | cm<sup>2</sup> |
|  | SLA_init | specific leaf area used to compute state variables at initialization | kg C/m<sup>2</sup> |
| ground area | pot_size | area of pot | cm<sup>2</sup> |
| growth scaling | root_to_shoot | root to shoot carbon allocation | mg/mg |
|  | leaf_to_stem | leaf to stem carbon allocation | mg/mg |

\*asterisk indicates parameters that were used for genomic prediction in this study

### Empirical evaluations

#### Baseline experiment

Experiments to characterize variation with respect to leaf growth and developmental rates across *B. rapa* morphotypes were performed at the University of Wyoming (Laramie, WY, USA). Plants were grown in the greenhouse (1800 μmol photons m^-2^ s^-1^ maximum PPFD, 20.75°C to 25.7°C /20.9°C to 21.3°C d/n, 26.5-73.8% relative humidity with an average of 47.1%) in the Fall of 2017. Three morphotypes of *B. rapa* were evaluated: five replicates of Chinese cabbage (CC), five replicates of vegetable turnip (VT) and sixteen replicates of oilseed (R500). Seeds of CC and VT were obtained from the Dutch Crop Genetic Resources Center (CGN) in Wageningen, VT (CGN10995) and CC (CGN06867) while seeds of R500, *B. rapa* subsp. *trilocularis* (Yellow Sarson), were part of a collection bulked at University of Wyoming in 2011. All seeds were planted in pots (500mL) filled with a soil mix (Miracle-Gro Moisture control Potting Mix (20% v/v), Marysville, OH, and Profile Porous Ceramic (PPC) Greens Grade (80% v/v), Buffalo Grove, IL) amended with 2 mL of Osmocote 18-6-12 fertilizer (Scotts, Marysville, OH) per pot. All plants were watered daily to maintain volumetric soil water content at 38% ± 5% (ECH_2_O; METER Group, Pullman WA). Individual leaf area were determined throughout the experimental period at least three times per week using Easy Leaf Area software (Easlon and Bloom, 2014) and its standard 4 cm^2^ red marker. Photographs were batch processed using the following parameter values: Leaf minimum Green RBG value (40-50 range), Leaf Green Ratio G/R (1.06), Leaf Green Ratio G/B (1.32), Scale minimum Red RGB value (0), Scale Red Ratio R/G & R/B (1.48), Scale area (4.0 cm^2^), Processing Speed (4), Minimum Leaf Size (0). Processed photos were inspected visually to ensure adequate green pixel coverage of each leaf. Leaf area data for these morphotypes may be found at https://github.com/DRWang3/leaf_model_TREES_paper.

#### Mild drought experiment

In the Spring of 2018 the mild drought experiment was conducted under greenhouse conditions at the University of Wyoming. Seeds of R500 and CC were planted using the same pot size and soil mix utilized in the baseline experiment of 2017. Plants were randomized and divided into three cohorts: a well-watered control group (WW) and two drought treatments (drought scenario 1 [D1] and drought scenario 2 [D2]) where drought was applied at two different developmental stages for each genotype. In total, the experimental set-up included 288 individual plants (54 D1 + 54 D2 + 36 WW per genotype) to account for necessary destructive sampling of whole plants for leaf water potential throughout the experimental period. In D1 and D2, water was completely withheld at the emergence of leaf four and emergence of leaf five, respectively, and plants were recovered just after emergence of leaf six and seven, respectively. Experimental overview presented in **Figure S2a**. Leaf area was measured approximately daily after the emergence of the third leaf using the Easy Leaf Area protocol described for the baseline experiment (**Dataset S1**). Projected canopy photos were taken at the emergence of the third leaf. Nine representative plants were tracked for leaf area in each drought treatment (total 9 replicates x 2 genotypes x 2 drought groups = 36 drought plants) while four plants were tracked in well-watered groups (total 4 replicates x 2 genotypes x 1 group = 8 well-watered plants). Leaf relative rate of expansion (RER) was computed following Granier and Tardieu 1999 as the local slope of the relationship between the natural logarithm of leaf area and thermal time.

Gas exchange and leaf water potential measurements, for computation of hydraulic conductance, were taken twice a day (predawn and midday) at regular timepoints throughout the experiment such that each genotype by treatment combination was evaluated twice during drought and once during the day after recovery (**Figure S2a, Dataset S1**). The leaf water potential assay required whole plant destructive sampling, so at each timepoint and time-of-day, six replicates *per* genotype by treatment (D1, D2) combination and two well-watered controls *per* genotype were measured. All measured plants were randomly selected and sample size was chosen according to successful experiments previously performed on *B. rapa* (Greenham et al., 2017; Guadagno et al., 2017). Pre-dawn and midday gas exchange measurements (Long and Bernacchi, 2003) were taken on the youngest expanded leaf. Photosynthetic rate and stomatal conductance were measured, using a portable gas exchange system (LI-C0R-6400XT; LI-COR Biosciences, Lincoln, Nebraska). Midday measurements (~10:00 a.m. to 1:00 p.m.) were taken in the conservatory where plants were growing, and environmental conditions in the cuvette matched ambient conditions in the greenhouse. Specifically, the cuvette settings were: flow rate, 300 μmol s^-1^; CO_2_, 400 μmol ^-1^ air; VPD, 1.1 ± 0.7 kPa, PAR (600 umol photon m^-2^ s^-1^ and block temperature set at 21°C, with cuvette fan on fast mode. Pre-dawn measurements (~3:00 a.m. to 5:00 a.m.) were taken with the same cuvette settings except in the dark and temperature set at 20°C; measurements at predawn were taken using a dim green light for visibility. For pre-dawn and midday leaf water potential measurements (Koide et al., 1989), plants were cut with a razor blade at the base of the stem and the main stem immediately inserted into a Scholander pressure chamber (model 600 Pressure Chamber Instrument; PMS Instrument, Albany, OR). Since the destructive sampling over the course of the experiment decreased the number of plants, individuals were randomized within their genotype by treatment trays after each timepoint.

Volumetric soil water content (VWC, %) (ECH_2_O; METER Group, Pullman WA) was monitored for all cohorts of plants to maintain the treatments for the duration of the experiment as shown in **Figure S2b**. This was measured daily in the evening for randomly selected pots within a genotype by treatment tray as well as at each TP for all pots of the plants being measured. A total of 589 VWC observations were taken in this manner over the course of 15 days. Soil VWC after recovery for all treatments was maintained at 38% ± 5% by keeping ~ 4cm of water at the bottom of each tray. For both empirical data collections (Fall 2017 and Spring 2018) all plants were harvested for above and below ground biomass evaluation. In the mild drought experiment, this was assessed at each timepoint in drought and at three additional days during recovery to assay biomass allocation over time. Above-ground tissue was cut at the soil level with a razor blade, weighed (fresh biomass), oven-dried for 10 days at 65°C and weighed again for dry biomass. The same procedure was applied to roots after harvest and manual gentle rinsing with DI water to eliminate soil and plant debris. Data collected from the mild drought experiment (leaf water potential, biomass, transpiration, leaf area data, and soil moisture) are found in **Dataset S1** as separate worksheets.

##### RIL leaf area analysis

Time-series leaf growth metrics (length and width of the 2^nd^ leaf) on the BraIRRi RIL population (Iniguez-Luy et al., 2009) were obtained from original observations under non-stressed growth conditions from a previous study also conducted at the University of Wyoming (Baker et al., 2015). To derive an estimated leaf area metric, leaves were assumed to have elliptical morphology, so area could be computed as

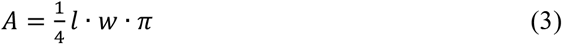

where *A* is leaf area, *l* is leaf length and *w* is leaf width. Thermal time was calculated based on the original study’s base temperature of 0.96°C and meteorological data provided by Baker *et al* (Baker et al., 2015). Hourly temperatures were 15.48-30.61^°^C with an average of 21.4^°^C, which encompassed the range experienced in the baseline experiment. Relative humidity ranged 35-48% with an average of 42%, which was a smaller range than in the baseline experiment.

##### Genotype parameterization

Genotype-specific parameters of logistic growth curves that model leaf area response to thermal time for the morphotypes (CC, VT, and R500) and the RIL population were estimated with a hierarchical Bayesian model with genotype and leaf levels. The genotype level was included because we were specifically interested in genotype-specific parameters and the leaf level was included because longitudinal measurements were taken on individual leaves, which may have similar or different behavior. Estimation was carried out using Gibbs Sampling implemented in the R package *rjags* (Plummer, 2013). Prior distributions for *K* and *r* were weakly informed from empirical means of the morphotypes and RIL population: dnorm(10, 25), dnorm(0.0007, 1e08), respectively, where the first value indicates the mean and the second indicates the precision. For *N_0_,* a prior of dnorm(0.01, 25) was selected based on theoretical knowledge that the initial value of a leaf growth curve should be very small (e.g., 0.01 cm^2^). After a burn-in period of 5,000 steps, four independent chains were sampled every 50^th^ sample and 1,000 steps were saved per genotype per parameter.

Univariate and multivariate scale reduction factors (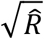 and 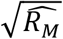) were evaluated to check for model convergence, ensuring that values were around 1.0 and never above 1.2 (Brooks and Gelman, 1998). Resultant posteriors of *K, N*_0_, and *r* per genotype were fit to gamma distributions using Maximum Likelihood Estimation implemented in the R package *fitdistrplus* (Delignette-Muller and Dutang, 2015) in order to generate gamma shape and rate parameters used by TREES as parameters for stochastic simulations.

##### Genomic prediction

Genomic prediction was performed on the BraIRRi RIL population (Iniguez-Luy et al., 2009) for the median, shape and rate estimates of each leaf growth parameter, *K, N*_0_, and *r*. A published genome-wide 1482 SNP dataset (Markelz et al., 2017) was utilized for 123 individuals that also had phenotypic data. After quality control, 1,481 SNPs with minor allele frequency higher than 0.01 were retained for genomic prediction. A total of eight approaches were analyzed, including a Genomic best linear unbiased prediction (GBLUP) model, Bayesian marker regression models, and machine learning models (reproducing kernel Hilbert spaces [RKHS] and random forest [RF]). The models differed in their assumptions of SNP effects and the counts of nonadditive genetic effects. Five-fold cross validation was performed using 50 replicates for each analysis and accuracy of genomic prediction was computed as the Pearson correlation between genomic breeding values and observed values.

#### GBLUP

The GBLUP model is:

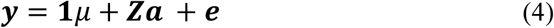

where **y** is the vector of the phenotype; **1** is the vector of ones; *μ* is the overall mean; ***a*** is vector of additive genetic effects accounted by SNPs; ***Z*** is the incidence matrix relating ***a*** to ***y; e*** is the vector of random residuals. It is assumed that 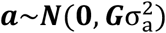 and 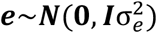. The G is the genomic relationship matrix constructed from SNPs followed method 2 of VanRaden (VanRaden, 2008); 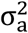 is the additive genetic variance explained by SNPs. The analyses using the GBLUP model was performed using the R package *sommer* (Covarrubias-Pazaran, 2016).

#### Bayesian marker regression models

The Bayesian marker regression models we applied were BayesA (Meuwissen et al., 2001), BayesB (Meuwissen et al., 2001), BayesC (Habier et al., 2011), Bayes Lasso (BL) (Park and Casella, 2008), and Bayes Ridge Regression (BRR) (Hoerl and Kennard, 1970). Models could be generalized as the following:

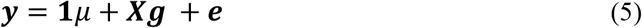

where ***y*, 1**, *μ* and ***e*** are the same as the GBLUP model; ***g*** is the vector of effects for SNPs; ***X*** is the genotype matrix, which has been centered and scaled for all genotyped individuals. Among four Bayesian marker regression models: BayesA and BL assume that SNP effects ***g*** have nonzero effects and the effect variances are drawn from scaled-t and double-exponential distributions respectively; BayesB assumes that SNP effects ***g*** are drawn from a two-component mixture of zero with probability n and a scaled-t distribution; BayesC assumes that SNP effects ***g*** are drawn from a two-component mixture of zero with probability n and a normal distribution; BRR assumes that SNP effects ***g*** are all drawn from a normal distribution with the same variance. Each Bayesian model was run as a single chain with a total length of 10,000 Markov chain samples, where the first 1,000 iterations were discarded as burn-in. Every 5th sample of the remaining 9,000 iterations was saved for the posterior analysis. The analyses using the Bayesian marker regression models were performed using the R package *BGLR* (Pérez and De Los Campos, 2014). Since the genetic background of the traits analyzed were unknown, default settings for hyper-parameters were used; for example, in the models BayesB and BayesC, the default rule is set to *xπ*_0_=0.5 and *p*_0_=10.

#### Reproducing Kernel Hilbert Spaces

Compared to the GBLUP model, the RKHS model used a Gaussian kernel function to measure the relationships among individuals (Gianola and Van Kaam, 2008). Due to the non-linearity of this Gaussian kernel function, RKHS could capture both additive and non-additive effects (Gianola and Van Kaam, 2008). The analyses using the RKHS model were performed using the R package *BGLR* (Pérez and De Los Campos, 2014). The bandwidth parameter of our analysis was set to 0.7 and setting for Markov chain was same as that for Bayesian marker regression models.

#### Random Forest

Implementations of RF have been shown to accurately capture epistatic effects (Motsinger-Reif et al., 2008; Michaelson et al., 2010; Picotti et al., 2013). Our RF analyses were carried out using *randomForest* package in R (Liaw and Wiener, 2002). The validation sets were predicted using a collection of regression trees with a subset of the total dataset that is bootstrapped over the total datasets. Then the predictions were averaged over all trees. The number of SNPs sampled at each split (mtry) was fine-tuned prior to the analysis using the package function tuneRF (ntreeTry = 500, stepFactor = 1.5, improve = 0.01). Random Forest was then applied with 10,000 trees (ntree).

##### Model exploration on an *in silico* population

The updated TREES model was examined on a simulated population of 123 individuals, which varied in the leaf growth module parameters *leafAreaMax, initialLeafSize, leafAreaRate, floweringTime,* and *SLA_low* and informed by the recombinant inbred line population, BraIRRi. All other parameters were fixed across the population in order to monitor the sensitivity of the model to growth and development constraints under varying environmental conditions. The *in silico* population was simulated under well-watered and water-limited scenarios. 40 non-flowering individuals were examined for ratios (water-limited to well-watered) for the following modeled traits: volumetric soil water content (%), leaf area index (cm^2^ cm^-2^) and canopy-level modeled transpiration (mmol H_2_O m^-2^ s^-1^), maximum potential transpiration (mmol H_2_O m^-2^ s^-1^), photosynthesis (umol CO_2_ m^-2^ s^-1^) and stomatal conductance (mol H_2_O m^-2^ s^-1^). Three individuals, selected based on *leafAreaMax* (at 25^th^, 50^th^ and 75^th^ quantile of population distribution of *leafAreaMax),* were further simulated using TREES’s stochastic mode to compare model results when informed by directly estimated parameters from data versus predicted by GP. 25 stochastic simulations were run per individual per treatment per parameterization method (25 simulations x 3 individuals x 2 treatments x 2 method [predicted versus direct] = 300 simulations). Parameter and drivers may be found at https://github.com/DRWang3/leaf_model_TREES_paper.

## RESULTS AND DISCUSSION

### Model development and parameterization

To adapt the process-based model TREES for studying short-lived crop species such as *B. rapa*, we replaced its original plant growth module with a new growth and development routine based on the sequential emergence and expansion of individual leaves. The general organization for this new module hinges on well-studied relationships between developmental rates and thermal time (Wilhelm and McMaster, 1995; Granier and Tardieu, 1998; Pantin et al., 2011) (**Figure 1 [step 1]**), which are governed by equations presented in **Appendix S1**. The 22 parameters of the sub-model describe leaf growth functions, temperature-dependent development, and resource allocation, each of which may be gathered from empirical evaluation (**Table 2**). To streamline a protocol for collecting input parameter data, we conducted a pilot experiment under non-stressed greenhouse growth conditions and characterized time-series growth traits of three *B. rapa* morphotypes of cultivated *B. rapa,* which differ dramatically in life-history traits and biomass allocation (oilseed [R500], vegetable turnip [VT], and chinese cabbage [CC]) (**Figure S3**). Of these cultivars, R500 was the only one to flower prior to the end of the experiment at 23,850 ^o^C hours (993.75 ^o^C days) (**Table S2**). CC exhibited high leaf emergence rate (i.e., a short phyllochron at 1027 ^o^C hours) and large final leaf size at 38.6 cm^2^, while VT and R500 displayed similar phyllochron (1980 and 1971 ^o^C hours, respectively), but R500 grew smaller leaves that took longer time to expand. The observed relative ranks were consistent with selection pressures that would have been necessary to drive the evolutionary divergence of these morphotypes; e.g., quickly emerging and expanding leaves and delayed transition to reproduction would have been favorable features for genotypes such as CC that are cultivated for consumption of vegetative organs. Of note, in all three genotypes the first two epicotylar leaves were consistently smaller at their maximum size than leaves that developed later (**Table S3a**).

**Figure 1.**
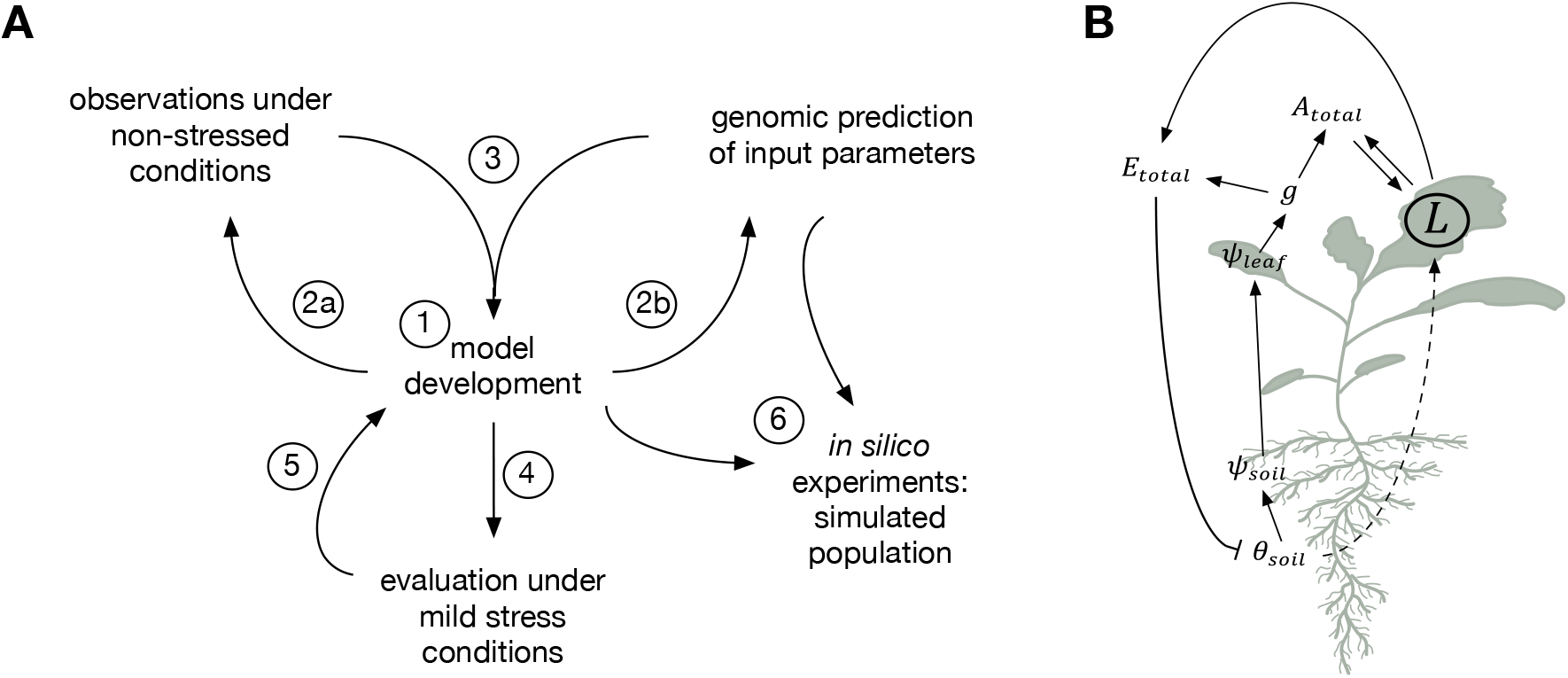
Study overview. **(A)** Summary of modeling, experimental, and analysis efforts carried out in this study. Stages: (1) A generalized sequential leaf growth sub-model is framed based on theoretical knowledge of dicot leaf development and integrated into the whole plant model, TREES. (2a-b) New model input parameters are tested for ease of direct measurement and on feasibility of indirect estimation using genomic prediction models. (3) Outcomes learned from 2a-b inform updates to model architecture. (4) Leaf growth and physiological traits are evaluated in response to mild drought. (5) Function for modulating leaf growth under water stress is incorporated to refine model. (6) The updated model is used to explore effects of growth and development variation on physiological response to mild water stress on a simulated population. Details, including plant material used at each stage, is found in **Table S1. (B)** A simplified systems diagram of TREES showing the high-level relationships between state variables of the original TREES model and the new leaf growth module (symbols: L, leaf growth module; A_total_, photosynthesis; E_total_, transpiration; g, stomatal conductance; ***ψ***_leaf_, leaf water potential; ***ψ***_soil_, soil water potential; ***θ***_soil_, soil water content). Dashed arrow represents the empirical function from Stage 5 of panel **A**.

**Table 2.** Comparison of accuracy across genomic prediction methods to predict leaf growth model parameters

| Trait | BayesA | BayesB | BayesC | BL | BRR | GBLUP | Random Forest | RKHS | Mean |
| --- | --- | --- | --- | --- | --- | --- | --- | --- | --- |
| *K* | 0.64 | 0.63 | 0.64 | 0.64 | 0.64 | 0.64 | 0.57 | **0.65** | 0.63 |
| *K_rate* | 0.6 | 0.6 | 0.6 | 0.6 | 0.6 | 0.6 | 0.55 | **0.64** | 0.6 |
| *K_shape* | 0.57 | 0.57 | 0.58 | 0.57 | 0.58 | 0.58 | 0.52 | **0.61** | 0.57 |
| *N<sub>o</sub>* | 0.42 | 0.42 | 0.42 | 0.42 | 0.42 | 0.42 | **0.45** | **0.46** | 0.43 |
| *N<sub>o_rate</sub>* | 0.5 | 0.49 | 0.5 | 0.5 | 0.5 | 0.5 | 0.48 | **0.56** | 0.5 |
| *N<sub>o_shape</sub>* | 0.5 | 0.5 | 0.51 | 0.5 | 0.5 | 0.51 | 0.5 | **0.57** | 0.51 |
| *r* | 0.31 | 0.31 | 0.31 | 0.31 | 0.31 | 0.31 | **0.34** | **0.35** | 0.32 |
| *r_rate* | 0.6 | 0.59 | 0.59 | 0.59 | 0.6 | 0.6 | 0.57 | **0.63** | 0.6 |
| *r_shape* | 0.59 | 0.59 | 0.59 | 0.59 | 0.59 | 0.59 | 0.56 | **0.63** | 0.59 |
| **Mean** | 0.52 | 0.52 | 0.53 | 0.52 | 0.53 | 0.53 | 0.51 | 0.57 | 0.53 |

Bold numbers indicate best-performing method(s) for each parameter. Accuracy is measured as the Pearson correlation between genomic breeding values and observed values and represent means from 50 replications of five-fold cross validation.

The central function that is computed at each 30-minute time-step in the updated TREES sub-model yields the relative rate of leaf growth under non-stressed conditions, equivalent to the logarithmic derivative of a leaf logistic growth function parameterized by *K, N_0_,* and *r* (Methods, Appendix S1). To derive genotype-specific inputs to the model, we estimated logistic leaf growth curve parameters for R500, VT, and CC off-line with the time-series leaf area data collected in the non-stressed baseline experiment (**Figure S4, Methods**). Estimated posterior distributions of the parameter *K*, the upper asymptote of the function, was the most differentiated across the three genotypes as compared to *N_0_* and *r*, reflections of initial leaf area and growth rate, respectively. Using medians of *K, N_0_*, and *r* for R500, CC, and VT to constrain the TREES model, we ran simulations for 1125 half-hour time-steps (~47 days) and evaluated longitudinal leaf growth. Results were consistent with expected leaf development in the three genotypes, with CC attaining high leaf area quickly and R500 growing small leaves slowly (**Figure S3** [bottom row]).

### Effects of trait uncertainty

Most process-based plant models are deterministic, i.e., one set of input parameters and environmental data yields a single time-series outcome. Such tools are useful to study system changes in response to time-varying environmental drivers, however, they are incapable of capturing within-genotype stochastic variation that would be useful for comparisons across genotypes over multiple simulation scenarios. In order to model a genotype’s phenotypic spectrum that is apparent even under experimental, non-stressed conditions, we asked whether we could inform process model inputs with parameter distributions in lieu of point estimates for leaf development. We fit gamma distributions to the posterior distributions derived from R500, CC, and VT for *K, N_0_*, and *r* parameters, which resulted in a shape and a rate estimate for each leaf growth parameter per genotype (**Figure S5, Table S2**). We then modified TREES accordingly to provide an option for internally sampling *K, N_0_*, and *r* out of distributions parameterized by rate and shape values (**Figure 1 [step 3], Appendix S1**). To explore the effects of general growth parameters and uncertainty in leaf-specific parameters on other traits modeled by TREES, we ran a series of stochastic simulations constrained for R500, VT, and CC growth and development. Simulated genotypes differed in leaf area index (LAI) (cm^2^ cm^-2^) (**Figure S6a-b**), which was expected due to its direct relationship with leaf area, but they also differed in whole plant hydraulic conductance (mmol m^-2^ s^-1^ MPa^-1^) (**Figure S6c-d**) and transpiration traits (mmol m^-2^ s^-1^) (**Figure S7**). Leaf water potential (MPa) was less differentiated overall but diverged in the latter part of the simulations, especially between CC and the other two genotypes (**Figure S6e-f**). Modeling outcomes suggested that leafy morphotypes such as CC may be more vulnerable to water stress than oeliferous types such as R500 (**Figure S7**), unless other intrinsic traits (e.g., those related to gas exchange) were concurrently divergent in these genotypes. Increased uncertainty of leaf parameters was observed to affect some traits more than others (**a, c, e** compared to **b, d, f** in **Figure S6–S7**); for example, while actual transpiration did not change appreciably, the time-series distributions of modeled maximum potential transpiration became considerably wider, giving rise to large ranges of hydraulic safety margin, i.e. the difference between transpiration and maximum potential transpiration (Sperry et al., 1998). Taken together, our results indicated that variation in growth parameters has the potential to give rise to variation in dynamic physiological responses. This was manifest despite only manipulating variation in growth parameters, highlighting how process-based models can capture internal interactions found inherently within plant systems.

### Genomic prediction of model parameters

Dicot leaf expansion, characterized by growth in two dimensions, is challenging to measure longitudinally in an automated fashion, with the exception of plants that form rosettes, such as *Arabidopsis thaliana* (Pantin et al., 2011). We therefore next asked whether genomic prediction can serve as an alternative to inform genotype-specific parameters in our process model, as part of assessing its potential scalability to large germplasm panels (**Figure 1 [step 2b**]). We utilized a previously collected genomic dataset (Markelz et al., 2017) (**Figure S8a**) and time-series morphometrics data (Baker et al., 2015) of the 2^nd^ leaf on an Recombinant Inbred Line (RIL) population, BraIRRi, derived from a cross between R500 and Imb211, a fast-cycling genotype (Iniguez-Luy et al., 2009). From the panel, we grew out two contrasting RILs to first evaluate development of the 2^nd^ leaf versus older and younger leaves. Size between the 2^nd^ leaf was observed to be more closely associated with sizes of the other leaves in these inbred lines than in the cultivars (**Table S3b**), likely due to the early-flowering nature of these RILs. This observation supported the use of the previously-collected data on 2^nd^ leaves to inform growth model parameters in TREES for this population. Genomic prediction of *K, N_1_*, and *r* posterior medians using a genomic best linear unbiased prediction model (GBLUP) were 0.64, 0.42, and 0.31 in accuracy, respectively. Notably, predictions of rate and shape gamma parameters were more accurate than predictions of the medians for *N_0_* (shape: 0.51, rate: 0.50) and *r* (shape: 0.59; rate: 0.60) (**Figure 2A**). In contrast, for K, whose median already had high prediction accuracy, accuracy decreased slightly (shape: 0.58, rate: 0.60). Simulations of parental leaf growth curves using 100 sets of *K, N_0_,* and *r* that were sampled from truncated gamma distributions (10^th^ −90^th^ quantiles) parameterized by predicted rate and shape values yielded outcomes that clearly discriminated the two genotypes; Imb211 curves overlapped with observed growth while R500 deviated from observations in the latter part of growth due to under-prediction of the *K* parameter distribution (**Figure 2B-E**).

**Figure 2.**
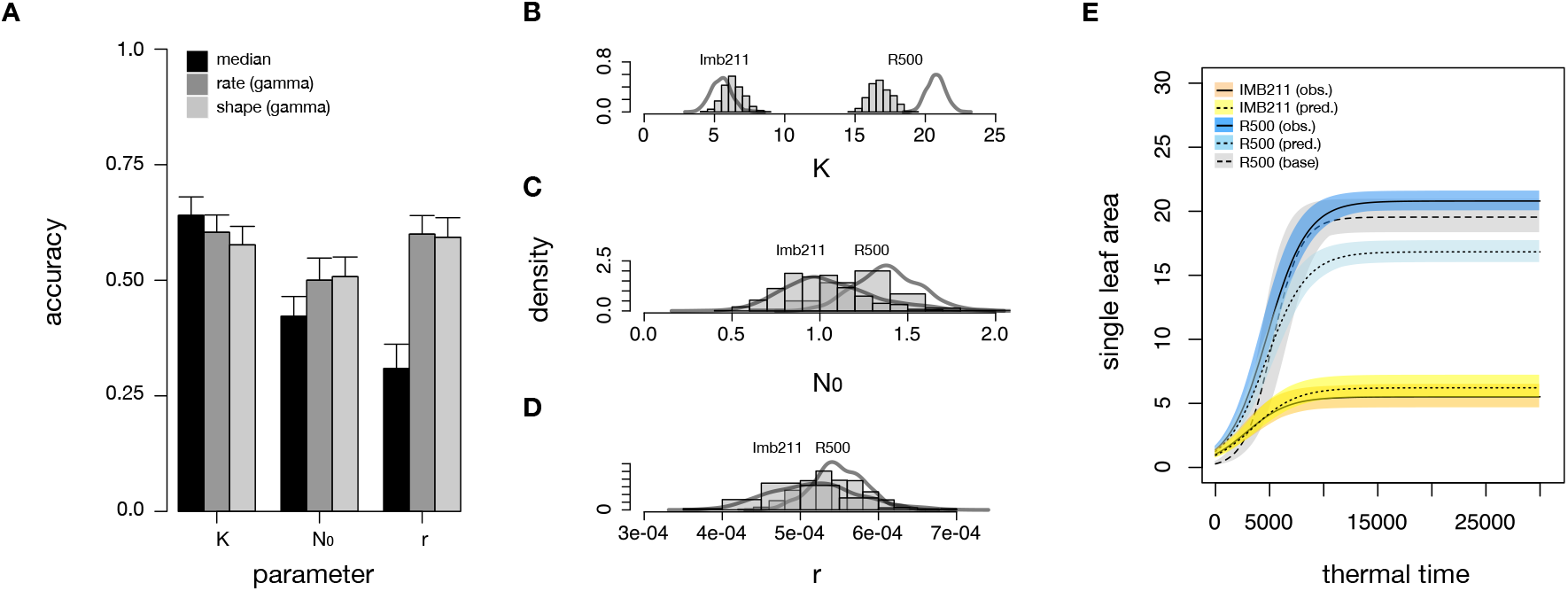
Genomic prediction of leaf growth parameters using a genomic best linear unbiased prediction (GBLUP) model. **(A)** Means from five-fold cross validation for predicting either single descriptors (medians) or distribution parameters of *K, N_0_*, and *r* posteriors in the Imb211 x R500 Recombinant Inbred Line (RIL) population. Bar shows 95% confidence interval of accuracy from 50 replicates of five-fold cross validation. Boxplots to show result distributions found in **Figure S8**. Accuracy is measured as the Pearson correlation between genomic breeding values and observed values. **(B-D)** Gamma distributions of leaf growth parameters (*K, N_0_*, and *r*) of parental lines R500 and Imb211 parameterized by rate and shape values either directly estimated (density curves) or predicted with ridge regression (histograms). Each density curve and histogram represent 500 samples. **(E)** Simulated leaf growth curves of R500 and Imb211 resulting from sampling from *K, N*_0_, and *r* truncate gamma distributions (10^th^-90^th^ quantiles) parameterized by rate and shape values either directly estimated (“obs.”) or predicted using genomic prediction (“pred.”). Also shown is simulated R500 leaf growth curves following the same sampling scheme using parameters directly estimated from the baseline experiment (“base”). Lines indicate the mean of the samples and shaded regions indicate range of 100 curves per class. Accuracy of genomic prediction was computed as the Pearson correlation between genomic breeding values and observed values.

We next compared genomic prediction performance across a suite of eight diverse models including GBLUP, four Bayesian marker regression models (BayesA, BayesB, BayesC, and Bayes Lasso [BL], and Bayes Ridge Regression [BRR]), and two machine learning models (Random Forest and Reproducing Kernel Hilbert Spaces [RKHS]). GBLUP and Bayesian regression models were selected because they are widely used in genomic prediction for plant and animal breeding, while the machine learning models were included for their potential to capture non-additive genetic effects. Genomic prediction accuracy is influenced by trait genetic architecture, which can be broadly defined in two dimensions: heritability (proportion of genetic variance to phenotypic variance) and complexity (number of genes controlling the phenotype). Overall, leaf model parameters showed moderate heritability (0.34-0.60 mean estimates across methods) and estimates were higher for the rate and shape parameters of *N_0_* and *r* as compared to their medians (**Table S4**). Prediction accuracies averaged across models ranged from 0.32 for *r* (median) to 0.63 for *K* (median) and largely reflected their heritability ranks (**Table 4, Figure S8b-l**). The five Bayesian marker regression models yielded similar prediction accuracies as GBLUP; these models perform similarly for traits of moderate complexity (Wang et al., 2018), suggesting moderate complexity of leaf growth parameters. To confirm the moderate complexity of leaf growth parameters, genome-wide association studies (McCarthy et al., 2008) could be carried out in future studies. Random Forest, which has been previously shown to uncover epistatic effects, was found here to outperform Bayesian marker regression models and GBLUP for *N_0_* (median) and *r* (median) but fared worse for other parameters. Overall, RKHS was favored, consistently achieving the most accurate prediction across all tested parameters, suggesting that non-additive genetic effects may play important roles for determining leaf growth traits. This was especially notable in predictions for rate and shape of *N_0_* where RKHS yielded accuracies 6% greater than other models. Interestingly, there was appreciable increase in accuracy for rate and shape parameters of *N_0_* and *r* distributions versus medians observed across all evaluated methods; this result provided additional support to modify TREES architecture to incorporate within-genotype uncertainty using rate and shape parameters (**Figure 1 [step 3**]). These empirical results supported the use of genomic prediction to inform genotype-specific leaf growth parameters as inputs into the updated TREES model.

### Leaf area modulation by mild drought

While empirical evaluation and genomic prediction lend critical information on growth progression under non-stressed conditions, the value of first principles-based process modeling lies in extending its predictive capabilities to novel environmental conditions, which may include changing environmental stressors such as transient water limitation. TREES is one of the few process-based models that couples rhizosphere and xylem hydraulics with carbon processes and has been shown to accurately predict water stress response across species such as aspen *(Populus tremuloides*), honey mesquite *(Prosopis glandulosa),* juniper *(Juniperus ashei),* and Texas persimmon *(Diospyros texana)* (Mackay et al., 2015; Tai et al., 2017; Johnson et al., 2018). Because there is evidence for the direct control of hydraulics on leaf expansion (Pantin et al., 2011; Caldeira et al., 2014), we set out to refine our growth model to capture leaf area modulation in response to water limitation in *B. rapa* by conducting a mild drought and recovery experiment using the genotypes CC and R500 (**Figure 1 [step 4]**). We imposed two staggered treatments of water-withholding (drought scenario 1 [D1] and drought scenario 2 [D2]), with their timings tailored according to leaf emergence events across the two genotypes (**Methods** and **Figure S2a** for experimental overview). Although water stress imposed was mild and only lasted between five and six days, droughted plants displayed visible loss of turgor prior to recovery (**Figure S9**) and mean biomass allocation had shifted to belowground under both drought treatments in each genotype (**Table S5**). Relative rate of leaf expansion (RER) showed a depression of RER during drought followed by a rapid increase after recovery (**Figures 3** and **S10**).

**Figure 3.**
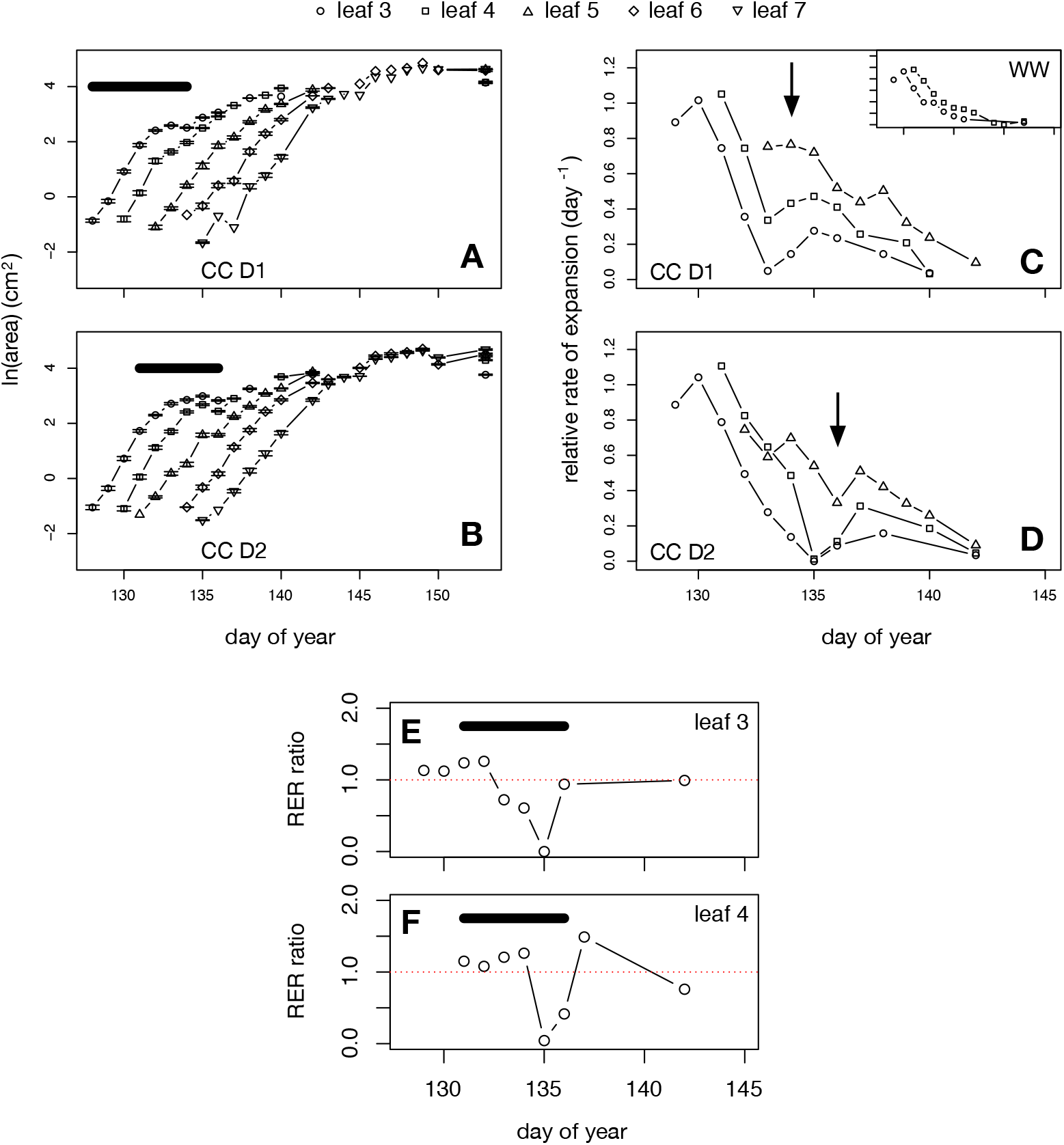
Leaf growth under mild drought. **(A-B)** Leaf area and **(C-D)** mean relative expansion rate of CC plants in response to mild water deficit and recovery. Error bars in **A-B** indicate standard error of the mean. Relative expansion rates in **C-D** were computed using means from A-B. **(E-F)** Ratio of the relative expansion rate (RER) in leaf 3 and leaf 4 of CC drought scenario 2 (D2) relative to CC well-watered treatment (WW); dotted line at 1.0 indicates equal RER in WW and D2. Arrows in **C-D** indicate onset of rewatering. Bars in **A-B, E-F** indicate periods of water limitation.

Using the data collected from the mild drought experiment, we sought to integrate functions into the TREES leaf growth module that could allow it to model leaf expansion under water-constrained conditions (**Figure 1 [step 5]**). Due to high variability and limited observations of hydraulic conductance (see Dataset S1 for leaf water potential and transpiration measurements), we opted instead to link the modulation of leaf area expansion to a measure of relative water status in the soil. This variable expressed a consistent signature across treatments (**Figure S2b**), and the empirical function related a soil water content ratio to a coefficient that modulates relative leaf area expansion rate (**Appendix S1**). To explore the updated whole plant model, we simulated the behavior of genotypes CC and R500 under their respective growth conditions beginning at the initiation of leaf 3 and compared modeled results to observed daily ‘slow’ traits (leaf area and volumetric soil water content) along with pre-dawn and midday ‘fast’ traits (leaf water potential and leaf-level measures of photosynthesis, stomatal conductance, and transpiration) (**Figures 4, S11–S13**). Overall, the model performed better for genotype CC than for R500 in nearly all traits, especially matching CC soil water content and leaf area progression well (RMSEs of 3.02% and 4.61 cm^2^). This likely indicates the need for better genotype-specific model parameterization for parameters that we had assumed species-level values due to lack of prior cultivar-level information; results suggest that those values may have been better suited for CC than R500. For leaf water potential, a challenging trait to monitor on tender herbaceous crops, noise found in the observed data in both genotypes led to a lack of strong differentiation between pre-dawn and midday values (**Figures S11c, S12e**) while in the model there was always a clear drop at midday due to the dependence of leaf water potential on hydraulics, leading to poorly matched modeled results for this trait. During periods of water deficit, relative expansion rate of simulated leaf growth (**Figure S13**) followed dynamics similar to what was observed (**Figures 3 and S10**). However, the empirical function we incorporated did not account for observed water-limited RER surpassing well-watered RER upon recovery, suggesting a knowledge gap in the mechanism of leaf expansion response to water recovery that may allow mildly droughted plants to ‘catch up’ to well-watered plants. Overall, the modeling link between soil water deficit and leaf expansion modulation serves as a functional placeholder until more data can be gathered to uncover how leaf RER responds to plant hydraulics.

**Figure 4.**
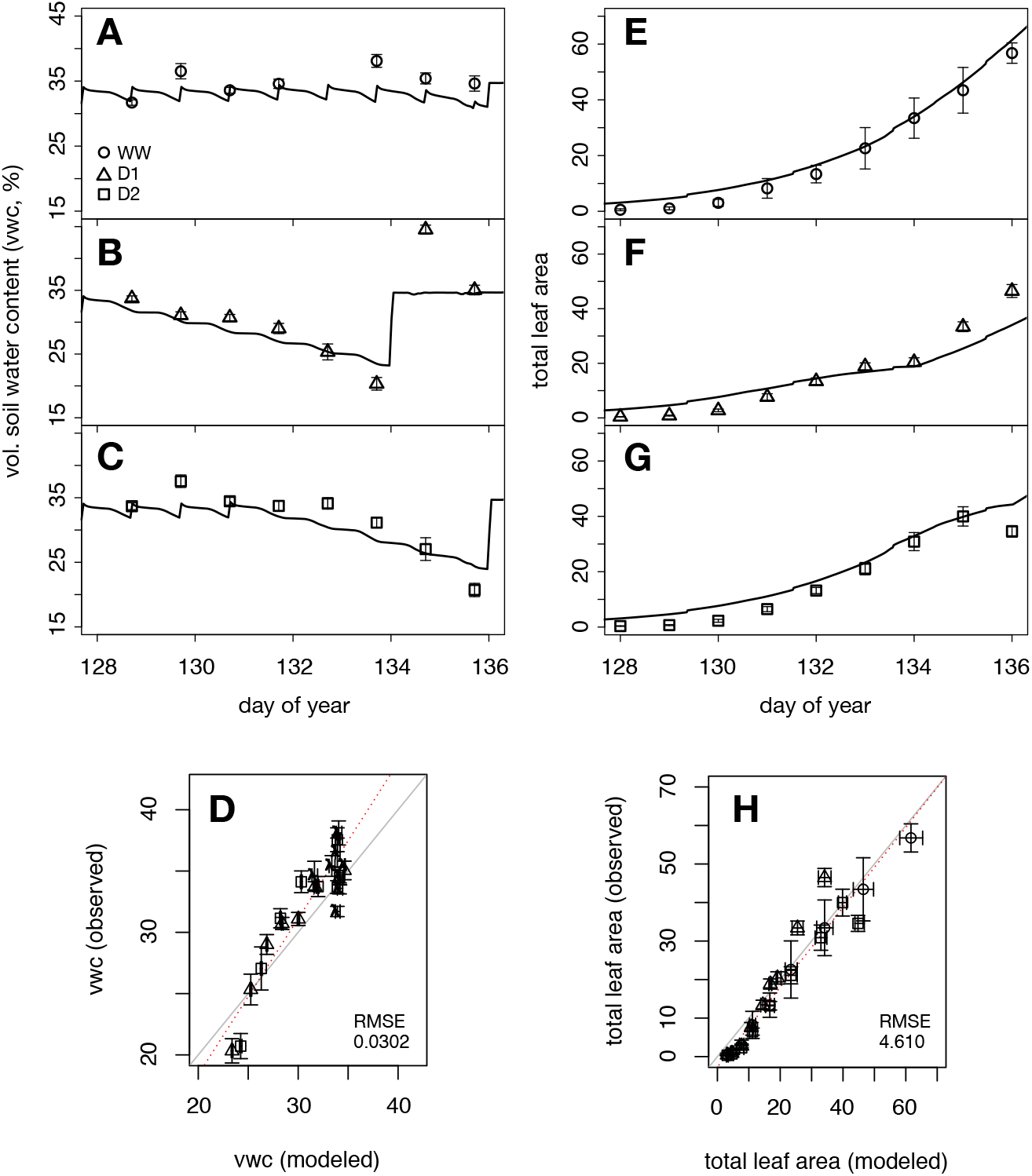
Confrontation of model with with observed soil water content and leaf area. TREES simulations parameterized with genotype CC leaf growth and forced with environmental data collected in the mild drought experiment. Time-series of soil water content (%) **(A-C)** and total leaf area (cm^2^) **(E-G)**. In one-to-one plots **(D, H)** solid lines represent the 1-to-1 line while dotted lines indicate best-fit. Observed points are means and error bars show their standard errors. Each modeled value shown in **D** and **H** is averaged from a 12-hour time interval corresponding to when the daily observations were made and bars indicate value range in the 12-hour intervals. Circle: WW (well-watered); triangle: D1 (drought scenario 1); square: D2 (drought scenario 2).

### A process-based modeling framework for decomposing G x E

While the new leaf growth module will benefit from future improved mechanistic understanding of drought response, its capacity to account for genotypic variation and within-genotype stochasticity afforded the opportunity to examine how process-based models might be useful for studying whole-plant genotype by environment interaction (G x E) (**Figure 1 [step 6]**). As a modelling exercise, we explored the behavior of a simulated population of *B. rapa* under 17-day well-watered and water-limited scenarios (**Figure S14**). Individuals (genotypes) of this *in silico* population varied for five leaf module parameters (*leafAreaMax, initialLeafSize, leafAreaRate, floweringTime, SLA_low)* associated with phenology and growth that were informed by a real *B. rapa* population, BraIRRi, and fixed for all other parameters. The water-limited scenario differed from the well-watered environment by having a six-day window of water deficit about halfway through the simulation (**Figure S15**).

We examined resulting trait ratios (water-limited to well-watered) of individuals in the population. Final ratios for leaf area index ranged from 0.787 and 0.931. This variation was largely associated with phenology differences in the set of genotypes that had transitioned to reproductive phase prior to the end of the simulation, with a correlation of −0.70 (p-val < 1.34e10^-13^) (**Figure S16**). Due to this association, we restricted the following examination to the 40 individuals that did not flower prior to the end of simulation. For volumetric soil water content, leaf area index, and transpiration traits, behavior was consistent across this population subset (**Figure 5A-B, Figures S17–S21**). In contrast, modeled canopy-level photosynthesis and canopy-level stomatal conductance exhibited behavior that diverged across this population subset (**Figure 5C-D, Figures S22–S23**). This divergent behavior was most apparent for stomatal conductance. After onset of recovery (days 7-13 in **Figure 5**), ratio ranges clearly encompassed the value 1.0, meaning that while some genotypes of the water-limited group had greater values after drought recovery than their well-watered counterparts, others had lesser values. The genotype rankings themselves were also dynamic, with some genotypes having >1.0 values at some timepoints in the recovery but <1.0 at others (**Figure 5D**).

**Figure 5.**
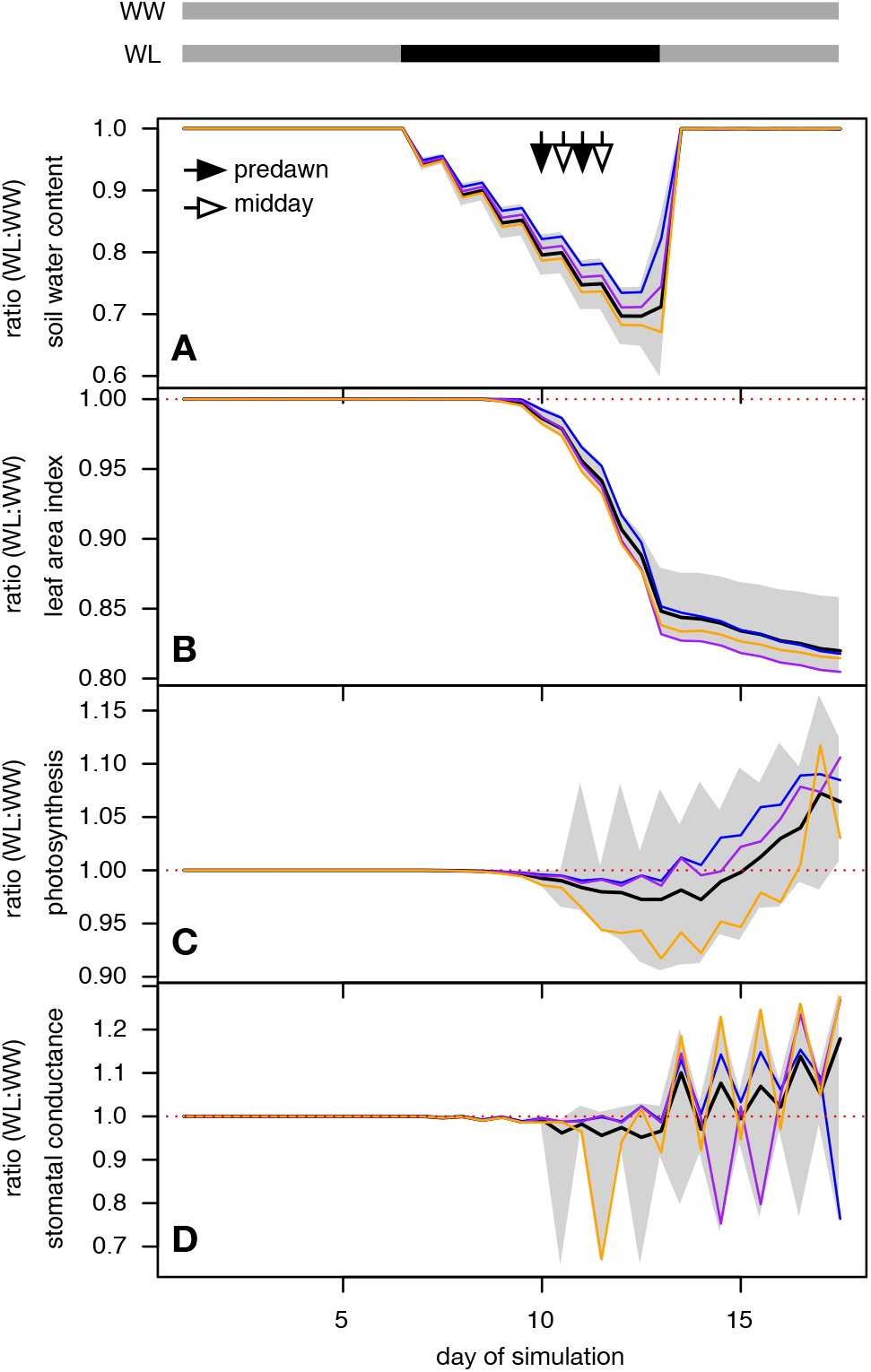
Behavior of a simulated *B. rapa* population under well-watered and water-limited scenarios. The 40 individuals of the *in silico* population that remained in vegetative phase for the entirety of the simulation period are compared for their behavior under well-watered (WW) and water-limited (WL) conditions (**Figure S15**). Watering regimes for each scenario are shown as gray and black bars at the top of the figure (gray indicates daily water input; black indicates no water input). Pre-dawn (5:00AM timestep) and midday (1:00PM timestep) ratios of WW to WL for **(A)** soil water content **(B)** leaf area index, **(B)** canopy-level photosynthesis, and **(C)** canopy-level stomatal conductance are shown here. Shaded gray region show the range of values across the 40 individuals; solid black curve depict the mean. Dotted line indicates 1.0, where WW and WL have the same value. Blue, purple, and orange lines depict three different individuals out of the 40.

These results were emergent properties of the system, in other words, outcomes of interacting model functions dependent on different combinations of variables integrated over time. While emergent properties are general phenomena of modeling complex systems, the specific example that we report here as emergent is a case of G x E, i.e. the differential response across genotypes to the environmental scenario (well-watered versus water-limited in this case). By exploring the updated model using an *in silico* population as opposed to a real population, we are able to safely conclude that the G x E we observed resulted from emergent properties of the model rather than other biological complexities characteristic of plant populations that are often challenging to account for (e.g., transgressive genetic variation). Additional stochastic simulations using three individuals contrasting in maximum leaf size (see **Methods**) supported genomic prediction as a means to parameterize growth in whole plant models for modeling across environmental scenarios (**Figure S24**). Taken together, results supported the updated model’s capacity to leverage genotypic-level differences in growth to drive intraspecific variability in response to environmental factors.

Previously, TREES utilized functions that depended on environmental variables *(e)* and/or population-level parameters *(p)*: generically *f(e), f(p), f(e,p)*. In the current study, additional layering of inter- and intra-genotypic variation allows for four supplemental relationship types constrained by genotype-level parameters *(g)*: *f(g), f(p,g), f(g,e), f(p,e,g)*. By formalizing the components that can give rise to emergent G x E, this update represents a theoretical advancement in the TREES model as it allows the dimension of time (and correlated environment) to interface with genetic variation. In the model, the relationship between genotype-specific leaf growth and temperature is an example of *f(g,e)* where *g* are the leaf growth parameters and *e* is thermal time; the new empirical modulation of relative leaf area by relative soil water content represents an *f(e)* function; computation of instantaneous leaf area index depends on functions that use *g* (leaf growth parameters) and *p* (parameter value for leaf angle fixed in the population for this case) (**Appendix S1**). Exploration of the *in silico* population provides evidence that the interaction of these and other new functions with original TREES routines integrated over time can give rise to whole plant G x E (**Figure 6**). Extending the modeling to real plant populations with proper genotype-level parameterization on all parts of the model (growth and development, nutrient cycling, carbon processes, and hydraulics) is needed to explore the predictive capacities of the model at-scale under stressed conditions. On the whole, this work suggests that the systematic identification, development, and testing of modeling components that can be refined by accounting for *g* may be key to utilizing process-based models for making predictions at-scale and identifying contributing elements underlying whole-plant G x E.

**Figure 6.**
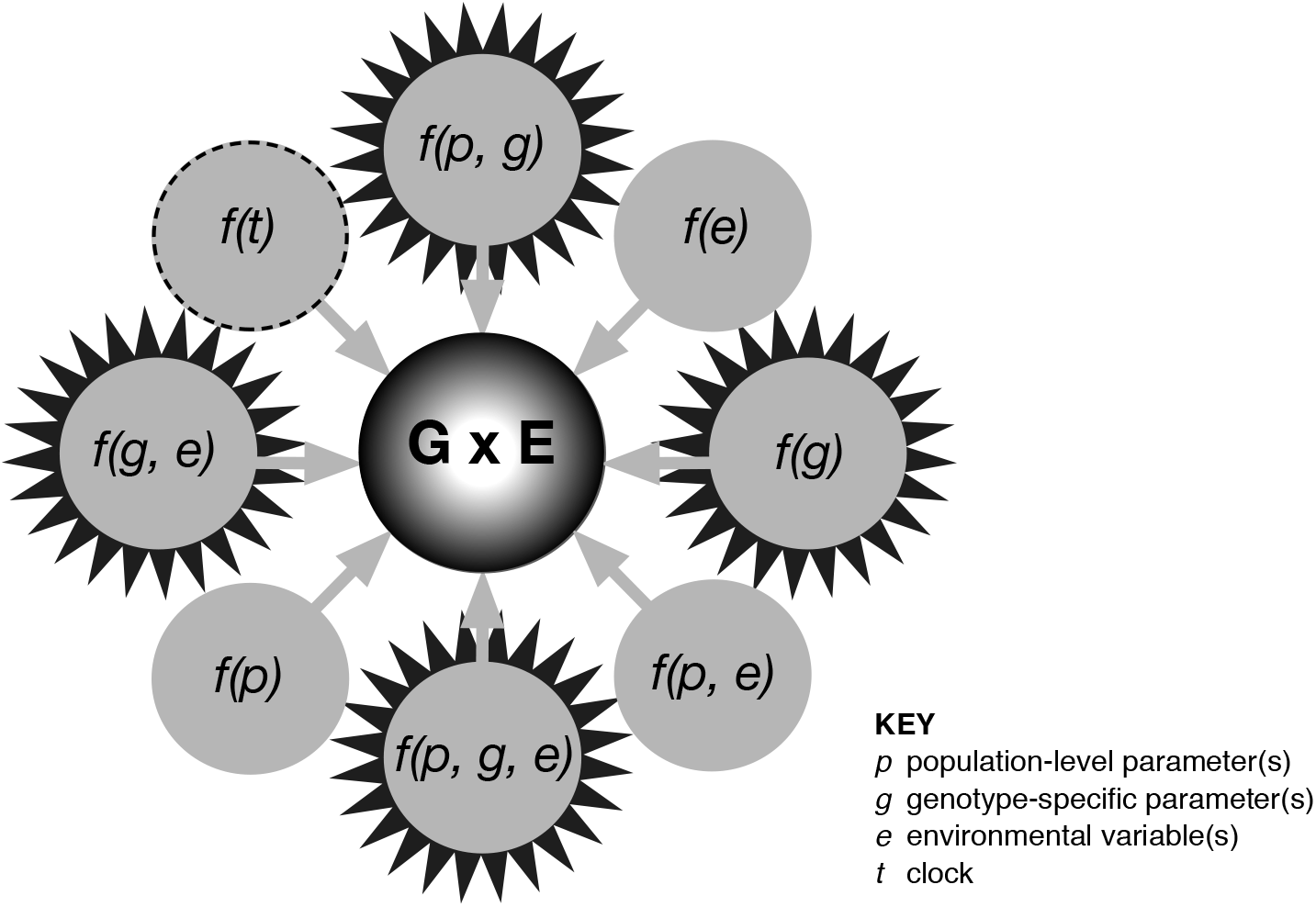
Decomposing genotype by environment interaction (G x E) with a biophysical framework. Genomics-informed process-based models are capable of formalizing the sub-processes that can give rise to observed genotype by environment interaction (G x E). G x E at the whole-plant level, an outcome of integration across multiple dimensions, is represented here by a ‘3D’ sphere. ‘2D’ circles are mathematical functions within the process model that are computed at each time-step of simulations. Not depicted are the routines that tie these various functions together; these may be thought of as single and double-headed arrows that link the 2D circles with each other. In this conceptual figure, the dimension of time extends straight out towards the reader, with multiple planes of circles stacked in layers on top of each other through time. Model state variables, which are carried over throughout a simulation from one time-step to the next, relate one layer to the next one above it. Starbursts mark model functions that utilize genotype-specific parameters, representing the parts of the model that can refined to account for genotypic variation. The *f(t)* circle with dashed outline are functions of the clock (i.e. circadian rhythm), another area for future model improvement.

## Conclusion

In this study, we introduced and tested the various components of a framework aimed at finding the modeling connections between genomics-informed design and environment-driven biophysics. First we overhauled the development module within TREES to adapt it for short-lived crop plants and modified model architecture to handle across- and within-genotype variation. Then we showed that genomic prediction offers a means to indirectly inform model input parameters for deterministic and stochastic simulations. Finally, we demonstrated the utility of a new empirical relationship in TREES to modulate the expansion of leaf growth under water-limited scenarios. The assemblage of experiments, analyses, and model updates laid out here represents a ‘staging area’ that supports a long-term goal of equipping plant process-based models with the ability to simulate large intraspecific populations. As technological advancements in plant phenotyping improve our ability to gather temporally-resolved data on large germplasm panels, models that can integrate natural genetic variation with physiological response will be key to harnessing this flood of information for application across both basic and translational domains.

## ACKNOWLEDGEMENTS

We thank Tim L. Setter for background resources that enabled initial conceptualization for the sequential leaf growth routine and James Berry for growth chamber access. We are grateful to Marta Szumski, Chris Nieters, Sarah Lemli Weber, Hunter Peterson, Rachel Shrode, Keegan Ferris, and Daniel Beverly for technical assistance. This study was funded by NSF-IOS #1547796.

## Supplementary file

**Figure S1.**
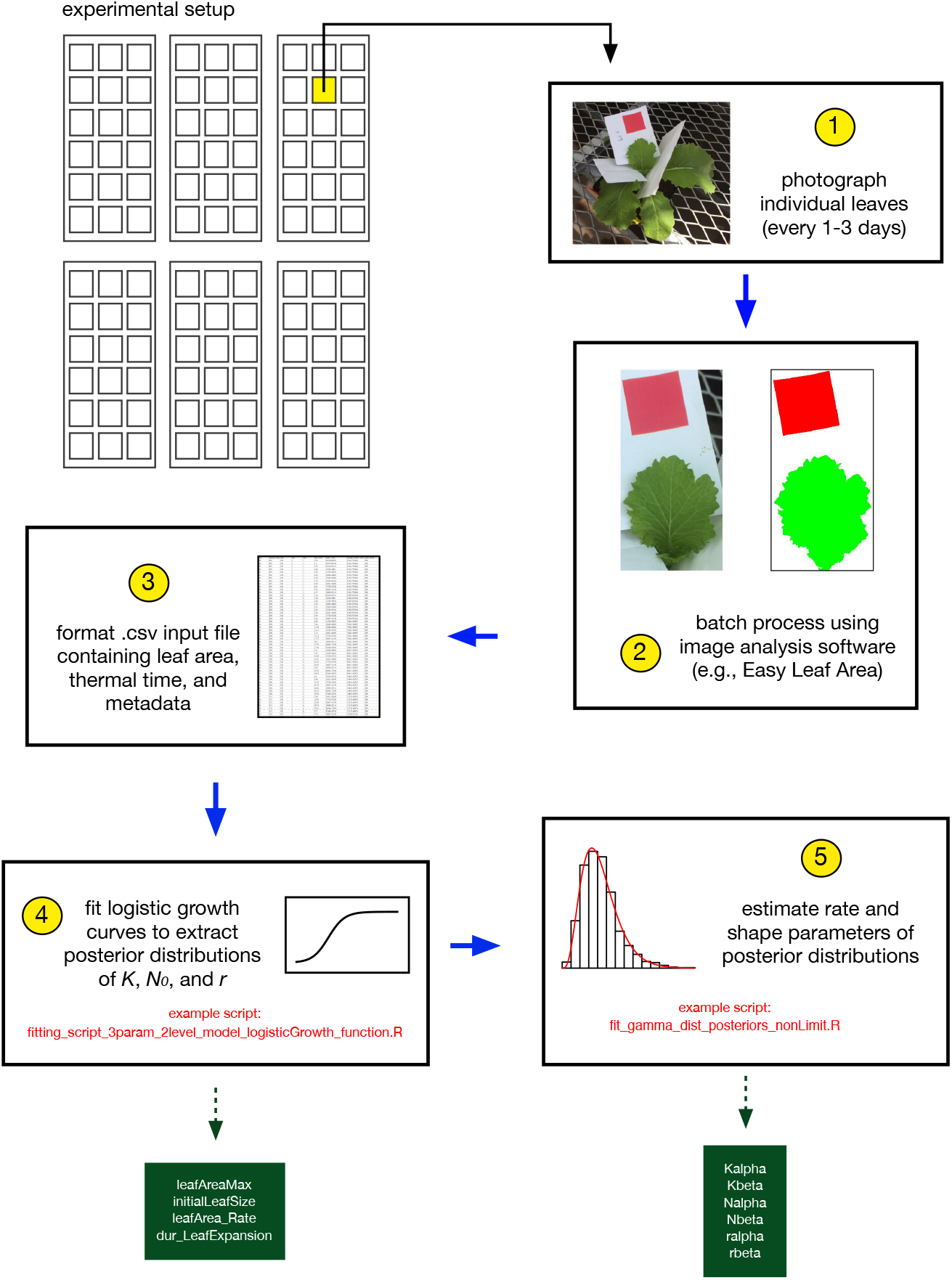
A pipeline to inform leaf growth input parameters of the TREES canopy growth submodel. (1) A representative subset of individuals per genotype are selected for leaf phenotyping. Photographs of leaf three or older (preferably leaf number four or five) are taken at high temporal density. (2) Images are processed to extract leaf area. (3) Data files formatted with to include thermal time. (4) Logistic curves are fit per genotype and resultant posterior medians are used for *leafAreaMax, initialLeafSize,* and *leafArea_Rate.* (5) Rate and shape parameters are estimated for posterior distributions and used for input parameters, *Kalpha, Kbeta, Nalpha, Nbeta, ralpha,* and *rbeta.* Scripts can be found at https://github.com/DRWang3/leaf_model_TREES_paper.

**Figure S2.**
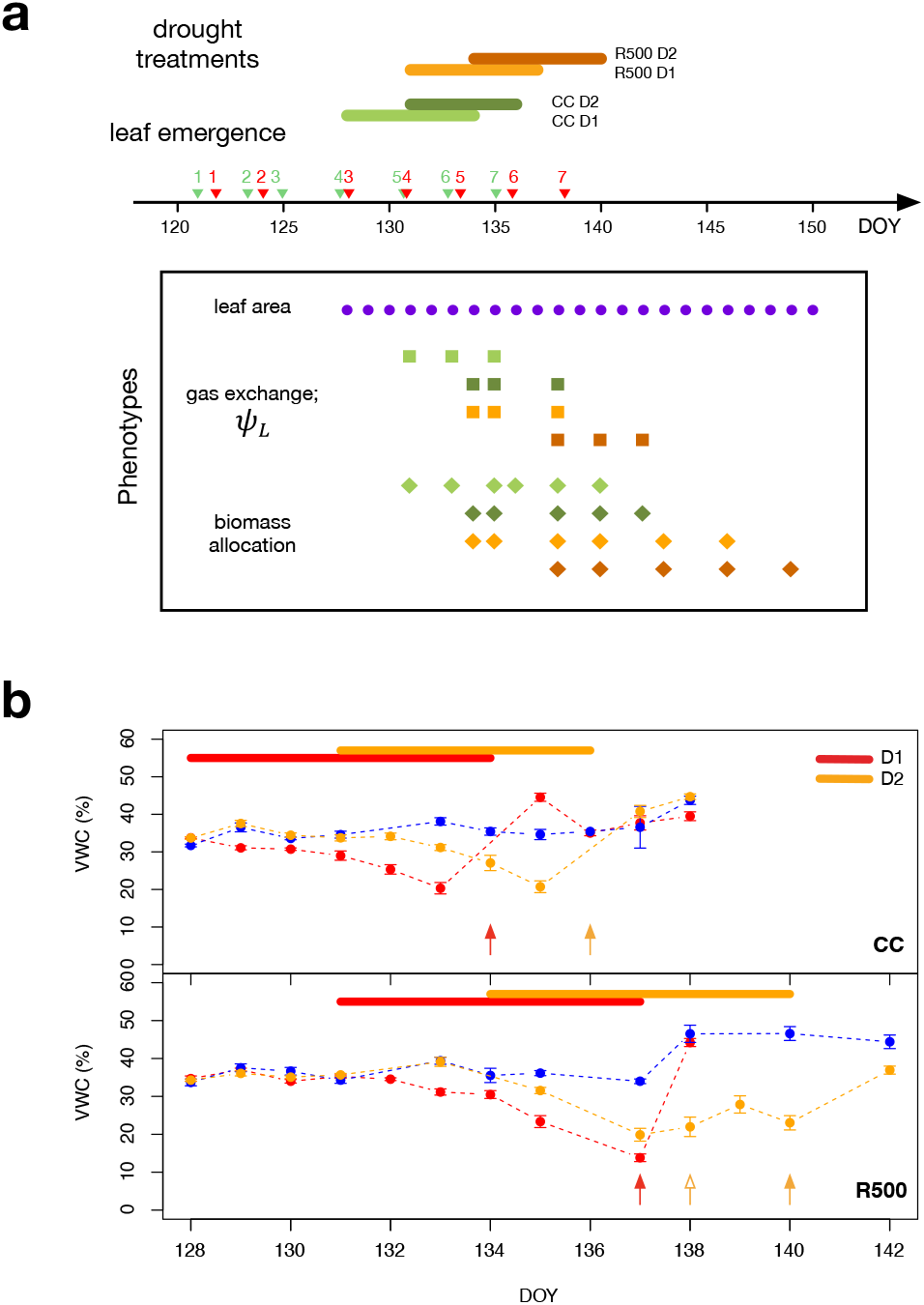
Evaluation of *B. rapa* genotypes under mild drought. **(a)** Experimental setup of the mild drought experiment. Overview of drought experiment showing the drought treatments (green and orange bars) for each genotype. Drought treatments were imposed according to developmental stage rather than clock time with leaf emergence events indicated in green and red triangles (green: CC; red: R500). Purple dots represent leaf area measurements using Easy Leaf Area. Whole plant destructive sampling for leaf water potential at pre-dawn and mid-day stages are marked in squares; gas exchange measurements for the same individuals were taken at those time-points. Above- and belowground biomass allocation was assayed throughout the experimental period, indicated by diamonds. **(b)** Volumetric soil water content (VWC, %) progression over the course of mild drought. Top panel for CC genotype and bottom panel for R500. Red and orange bars across the tops of each plot indicate drought duration for each treatment. Arrows with solid heads indicate the day drought was relieved. Open-headed orange arrow marks the day that a 5-minute bottom-watering was applied to the R500 D2 treatment prior to the end of drought because treated plants appeared excessively stressed from visual observation (e.g., portions of older leaves becoming dried). DOY: day of year (Julian Day).

**Figure S3.**
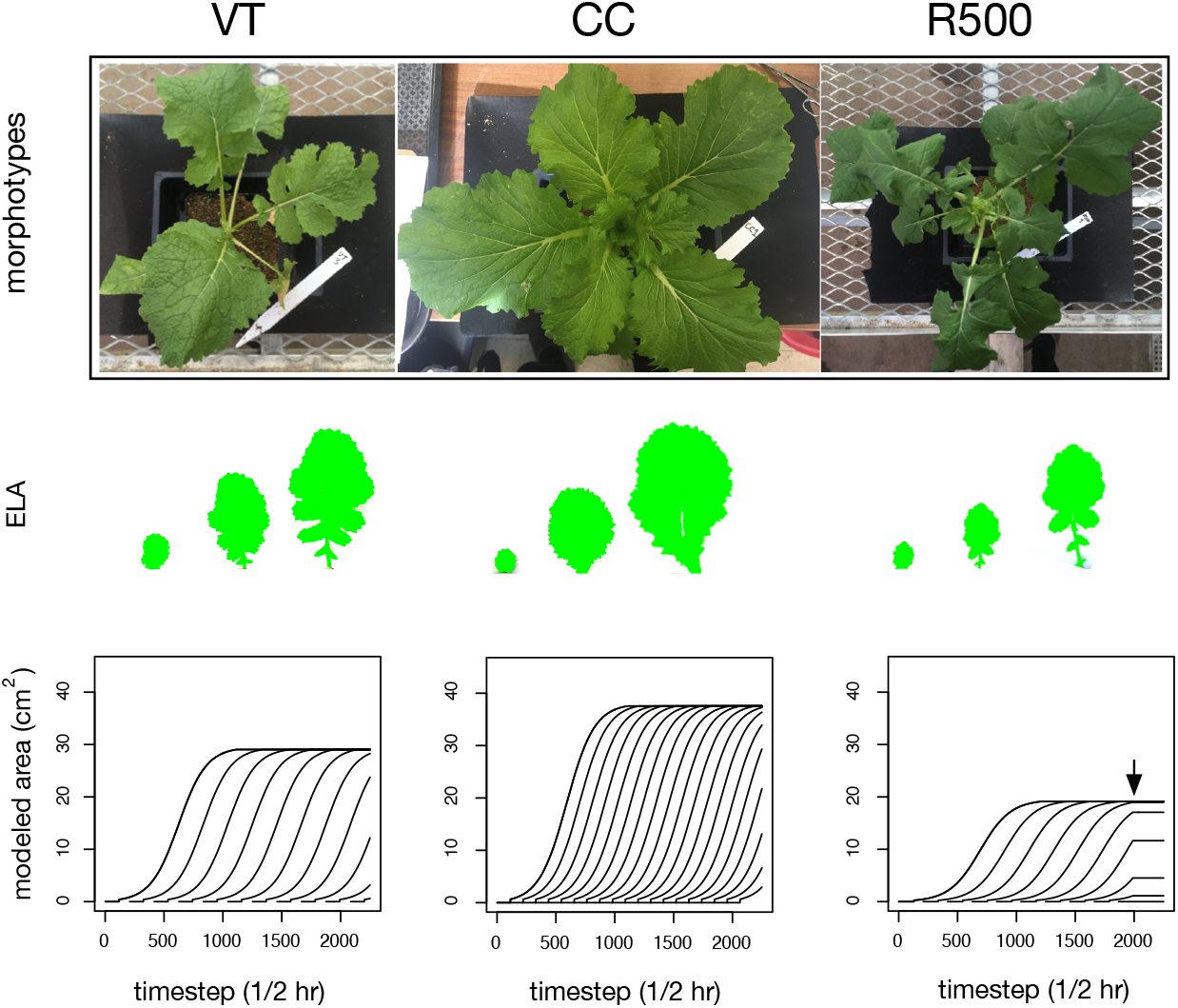
Vegetative leaf growth variation across *B. rapa* morphotpes. Baseline experiment evaluating genotypic differences in growth across three *B. rapa* morphotypes under non-stressed conditions. First row: photos of vegetable turnip (VT), Chinese cabbage (CC), an oilseed cultivar, R500, taken 53 days after sowing. Second row: Processed Easy Leaf Area (ELA) images from leaf five in a representative individual at three timepoints throughout leaf expansion. Third row: TREES simulations of morphotype-specific leaf growth under non-stressed conditions. Arrow indicates a transition to the reproductive phase for R500 in the model.

**Figure S4.**
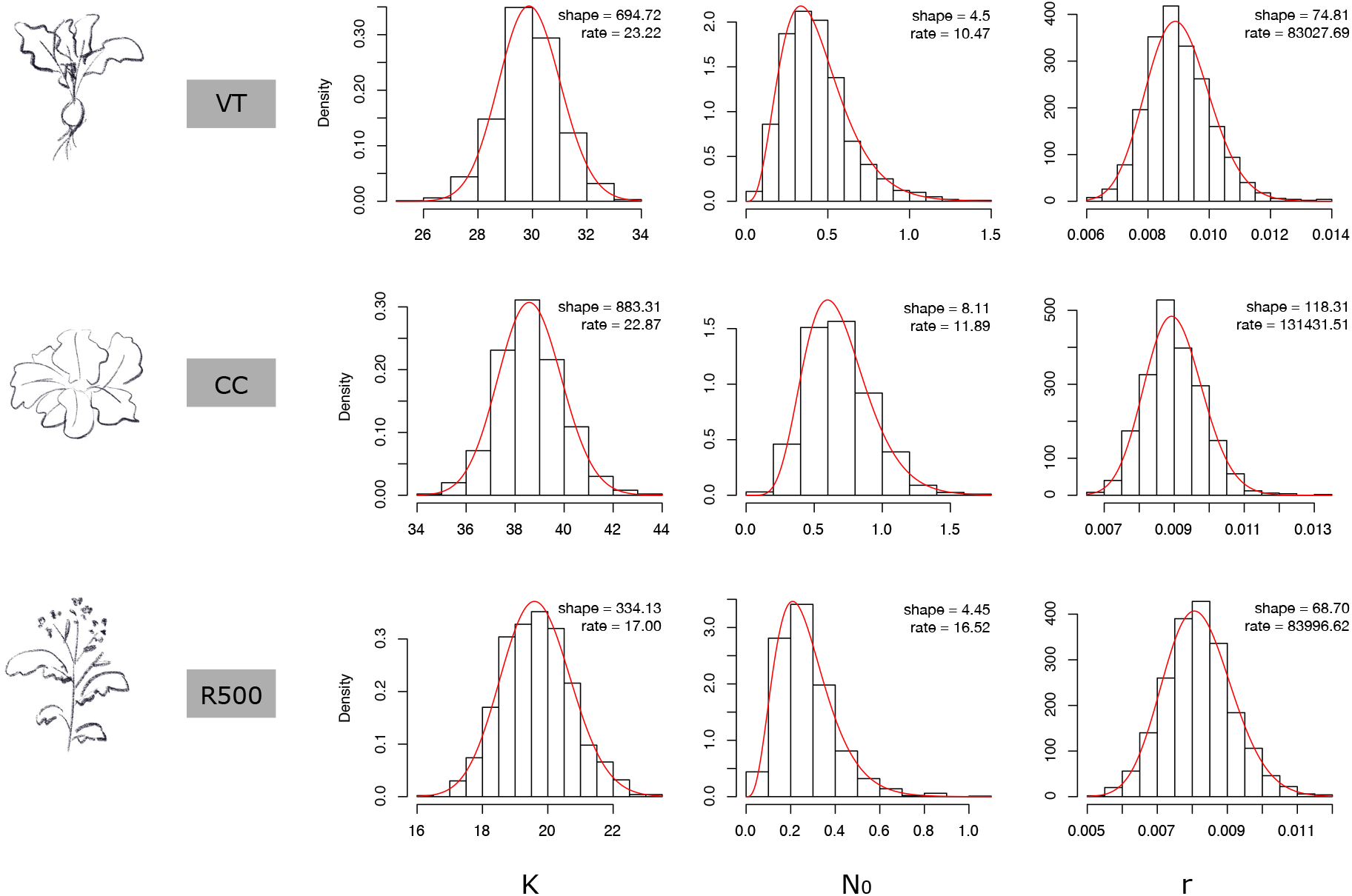
Posterior parameter distributions of leaf growth curves across *B. rapa* morphotypes. Histograms depict posterior distributions from Bayesian parameter estimation of *K, N_0_*, and *r* parameters using two-level hierarchical models (levels: genotype and leaf). Red lines represent fitted gamma distribution. Shape (α) and rate (β) parameter estimates are indicated within each figure. The number of individual plants per genotype were five replicates for CC, five replicates for VT, and 16 replicates for R500. Time-series area measurements for leaf number three and greater were used for parameter estimation. The script for estimating these parameter posterior distributions is provided at https://github.com/DRWang3/leaf_model_TREES_paper.

**Figure S5.**
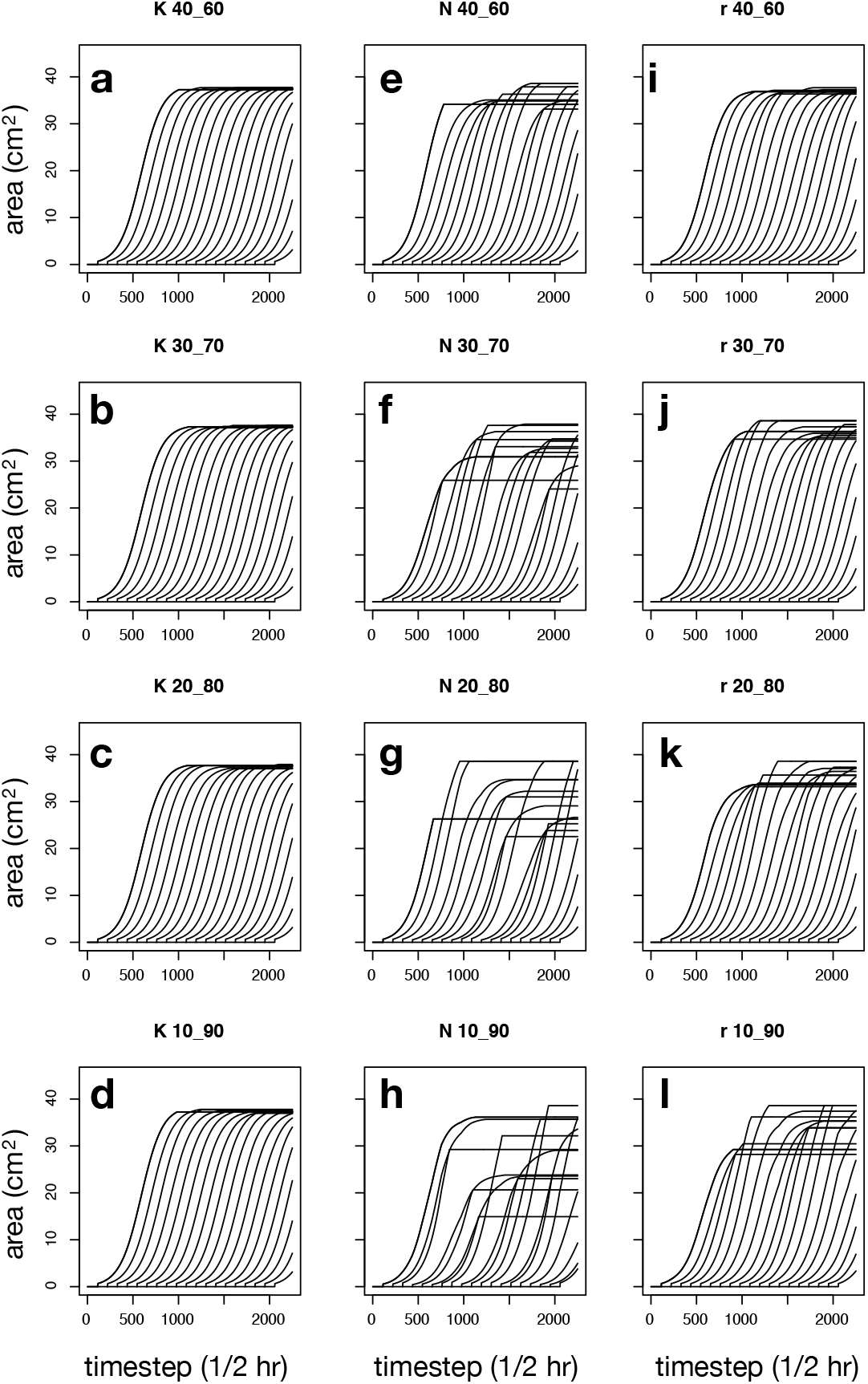
Stochastic leaf growth curves for CC. Growth of each individual leaf is parameterized by different *K, N_0_*, and *r* values sampled from truncated gamma distributions (gamma distribution parameters found in **Figure S3**). **(d-g)** Randomly sampled *K* with fixed *N_0_* and *r* values. **(h-k)** Randomly sampled *N_0_* with fixed *K* and *r* values. **(l-o)** Randomly sampled *r* with fixed *K* and *N_0_* values. Truncated ranges are indicated in plot titles (e.g., “40_60” indicates sampling the distribution within the 40^th^ – 60^th^ quantile range).

**Figure S6.**
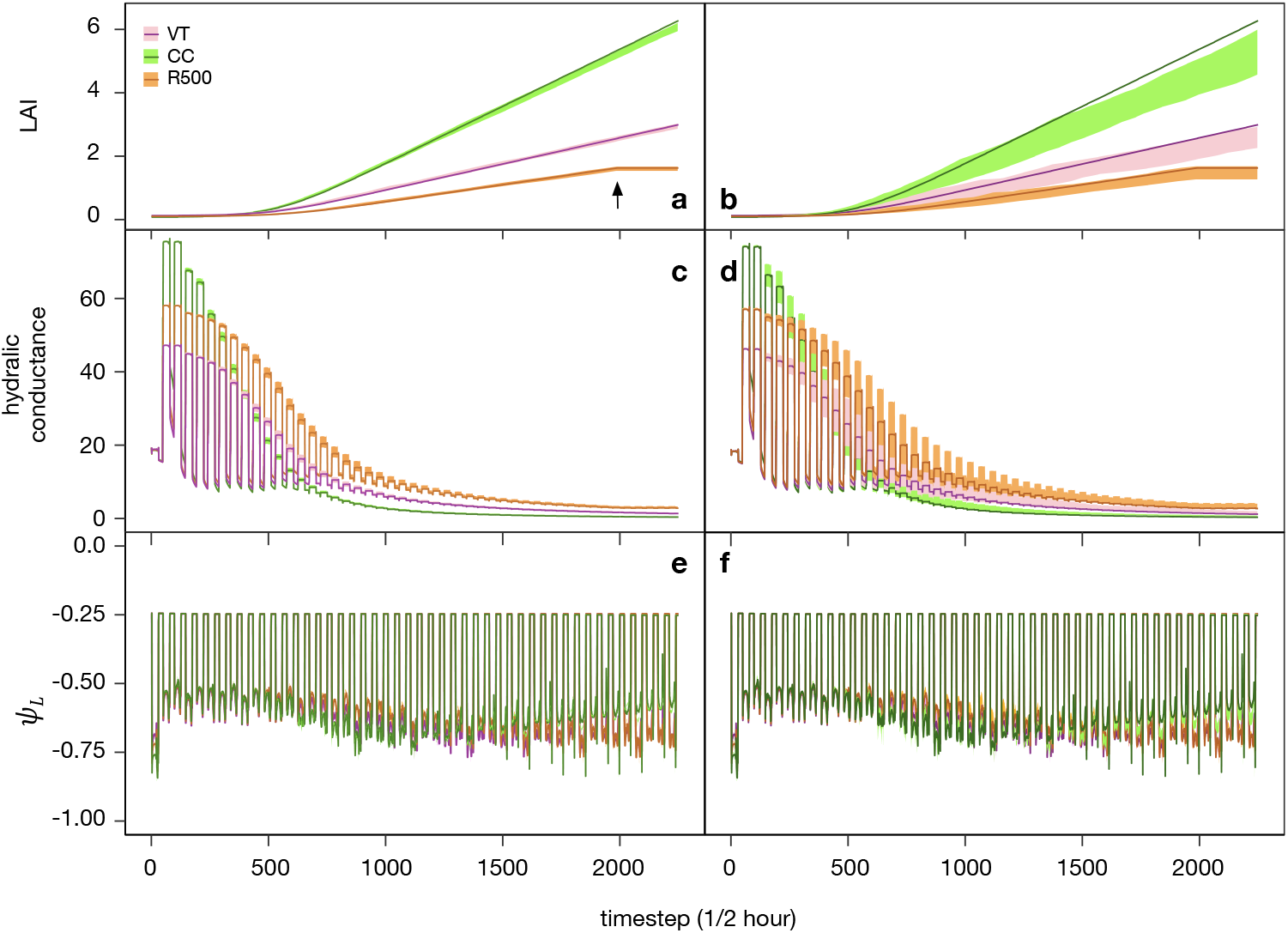
Using genotype-specific stochastic parameters to inform whole plant simulations. Time-courses (~47 days) of *Brassica rapa* using leaf growth parameter distributions derived from genotypes VT, CC, and R500 truncated between the 40th and 60th quantiles. Shown here are differences in **(a-b)** Leaf area index (LAI, cm^2^ cm^-2^), **(c-d)** whole plant hydraulic conductance (mmol H2O m^-2^ s^-1^ Mpa^-1^), and **(e-f)** leaf water potential (MPa) driven only by differences in growth parameters. All other input parameters were set as equal. Plots summarize 50 simulations per genotype (line: simulation using fixed growth parameters; shade: 50 simulations using truncated parameter distributions). **(a, c, e)** show results from sampling from narrow distributions (range 40th to 60th quantile) and panels **(b, d, f)** displays results from sampling from wider parameter distributions (range 10th to 90th quantile).

**Figure S7.**
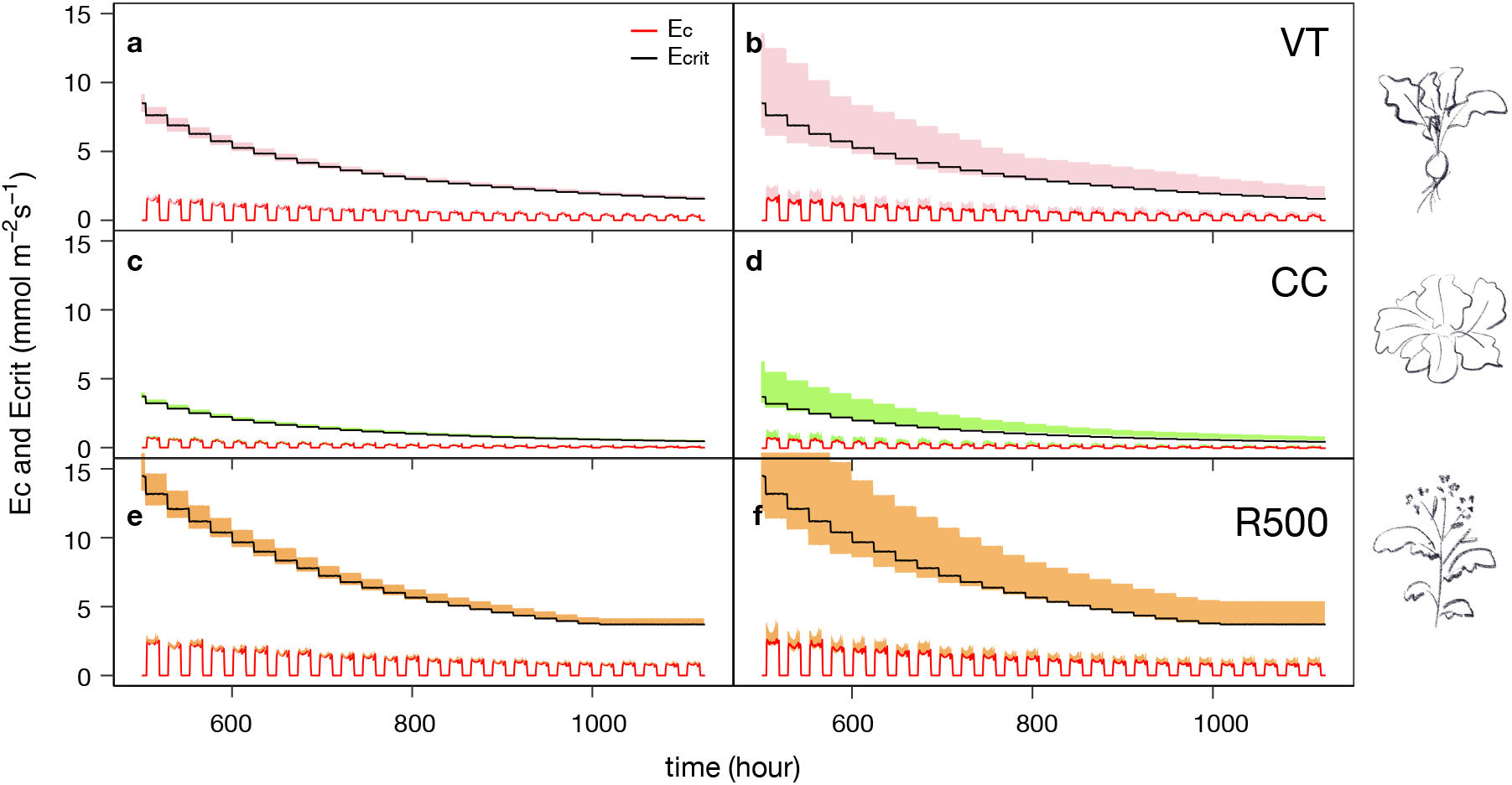
Transpiration outcomes in simulations with stochastic leaf growth. E_c_ (modeled transpiration, mmol H2O m^-2^ s^-1^) and E_crit_ (modeled transpiration potential, mmol H2O m^-2^ s^-1^) time-courses in the latter part of the simulation (hour 500 to hour 1125) for **(a-b)** VT **(c-d)** CC and **(e-f)** R500. Red line: deterministic outcome for E_c_; black line: deterministic outcome for E_crit_. Pink, green, and orange shaded regions indicate the ranges of outcomes for E_c_ and E_crit_, respectively, that resulted from 50 simulations each for VT, CC, and R500. Parameters were sampled from truncated distributions in the 40th-60th quantile range **(a, c, e)** or 10th-90th quantile range **(b, d, f)**. Illustrations on the right depict morphological differences across VT, CC, and R500.

**Figure S8.**
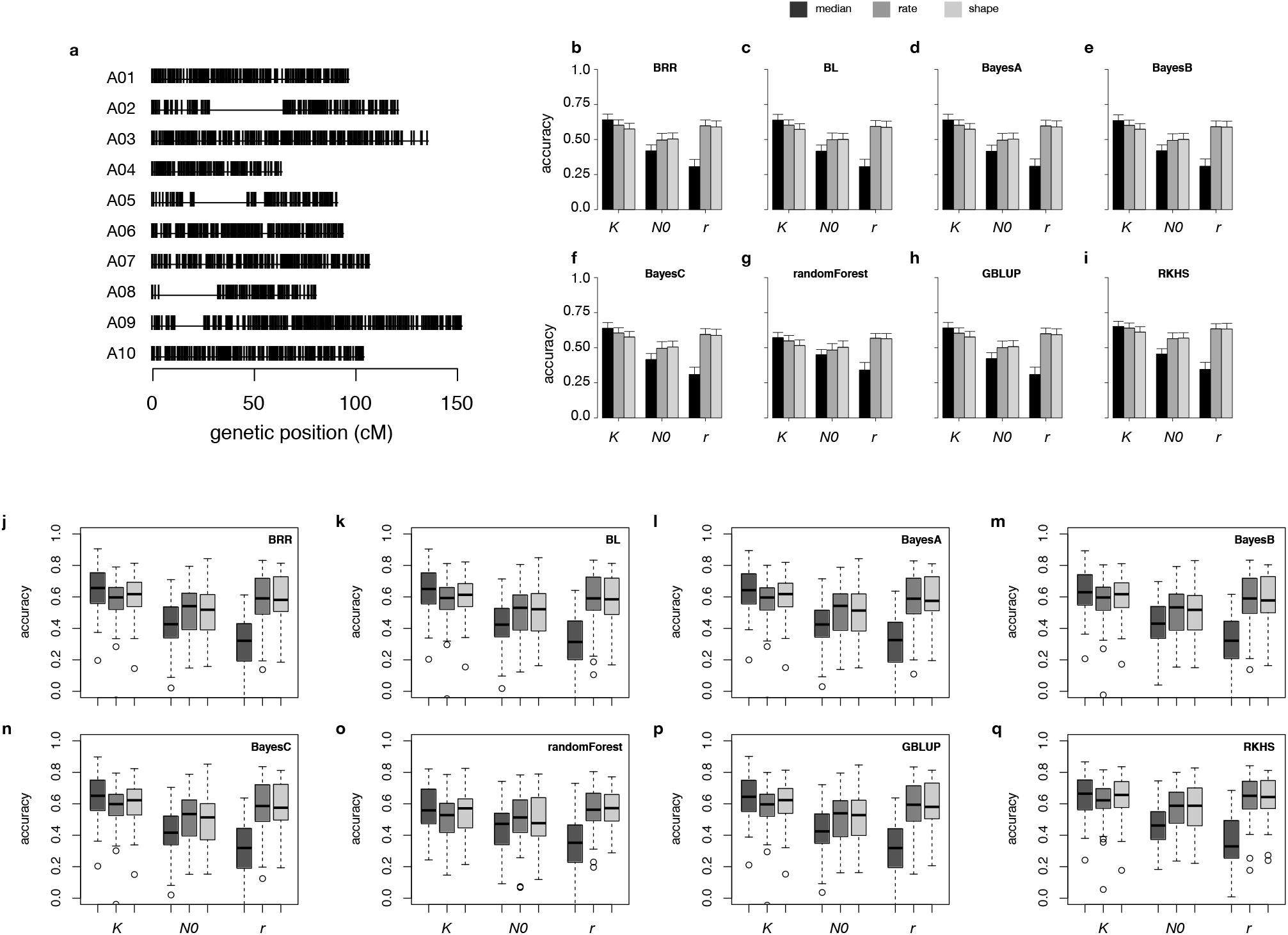
Genomic prediction of leaf growth model parameters. Eight genomic prediction approaches are compared for predicting the median, rate parameter, and shape parameter of *K, N_0_*, and *r* leaf growth parameters in a bi-parental recombinant inbred line population. **(a)** Distribution of 1481 SNP markers across ten chromosomes used for genomic prediction. Results from Bayesian marker regression models **(b-f)**, a genomic best linear unbiased prediction model **(h)** and machine learning models **(g, i)**. For accuracy values, see **Table 2**. Error bars: 95% confidence interval estimated from 50 replicates of five-fold cross validation. Boxplots found in **(j-q)** are generated from the same 50 cross-validation replicates in **(b-i)** and are included here to show distributions. Colors used in **(j-q)** refer to the same legend as for **(b-i)**. Accuracy of genomic prediction was computed as the Pearson correlation between genomic breeding values and observed values.

**Figure S9.**
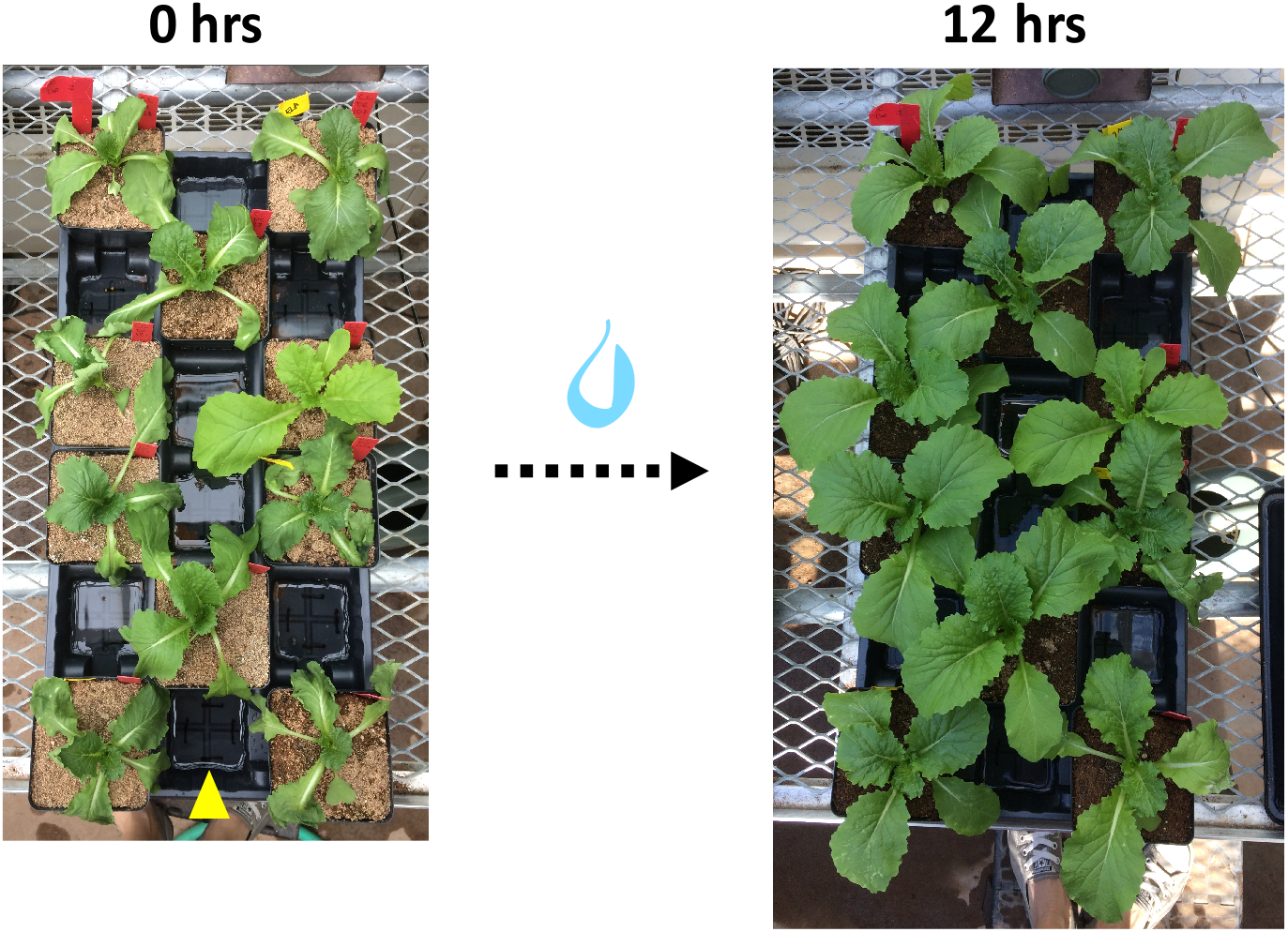
Recovery of droughted plants. After a period of drought synchronized with development, drought was relieved with bottom-watering. Shown here is one tray of CC D2 plants at the onset of recovery (0 hours; left photo) and the same tray after overnight recovery (12 hours; right photo). Yellow arrowhead indicates the bottom water level added at time of recovery (~4cm in height).

**Figure S10.**
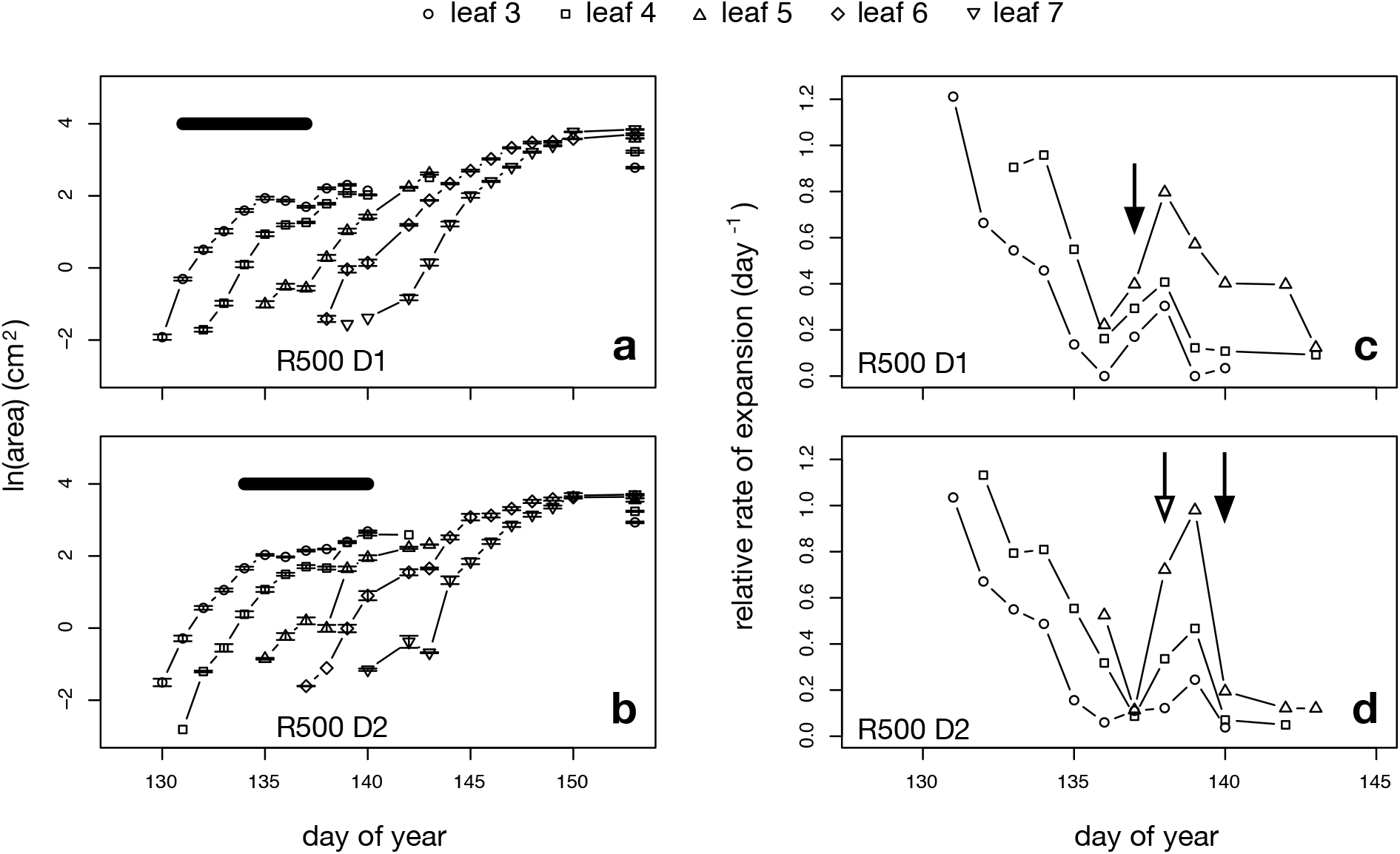
R500 leaf growth under mild drought. **(a-b)** Leaf area and **(c-d)** mean relative expansion rate (day^-1^) in response to mild water deficit and recovery. Error bars in **a-b** indicate standard error of the mean. Relative expansion rates were computed using means in **a-b**. Bars in **a-b** indicate periods of water limitation. Arrows in c-d indicate onset of re-watering.

**Figure S11.**
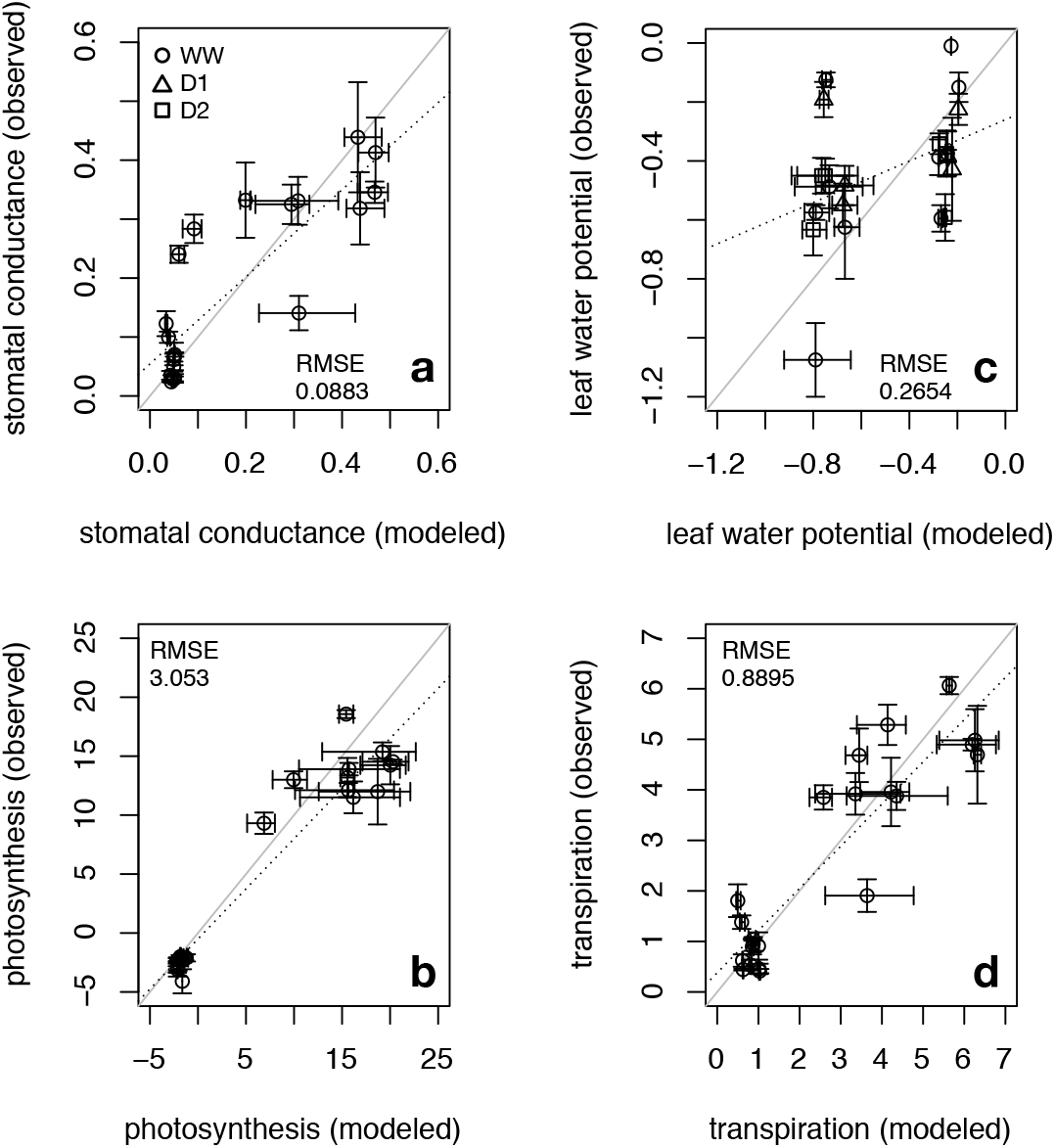
Simulated versus observed traits in genotype CC. TREES simulations parameterized with genotype CC leaf growth and forced with environmental data collected in the mild drought experiment. Traits of interest are **(a)** stomatal conductance (mol m^-2^ s^-1^), **(b)** leaf-level photosynthetic rate (umol CO_2_ m^-2^ s^-1^), **(c)** leaf water potential (Mpa), and **(d)** transpiration (mmol H_2_O m^-2^ s^-1^). Gray line: 1-to-1 line; dotted black line: best-fit line. Observed points are means and error bars indicate standard error of the mean. Each modeled value shown is averaged for the time interval corresponding to when gas exchange and leaf water potential measurements took place and bars indicate modeled value range. Circle: WW (well-watered); triangle: D1 (drought scenario 1); square: D2 (drought scenario 2).

**Figure S12.**
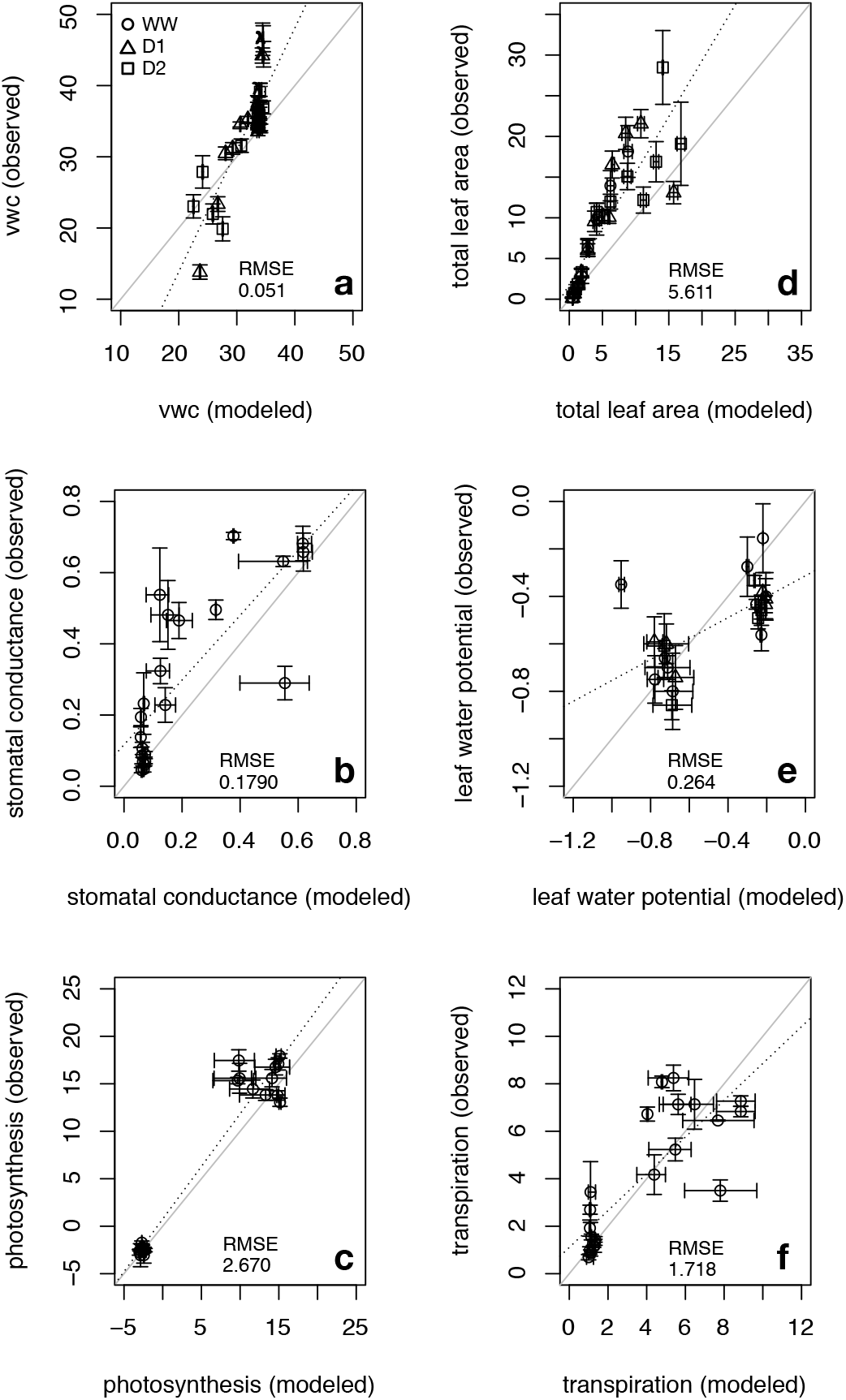
Simulated versus observed traits in genotype R500. TREES simulations parameterized with genotype R500 leaf growth and forced with environmental data collected in the mild drought experiment. Gray line: 1-to-1 line; dotted black line: best-fit line. Observed points are means and error bars indicate standard error of the mean. Each modeled value shown for volumetric soil water content (%) **(a)** and leaf area (cm^2^) **(b)** are averaged from a 12-hour time interval corresponding to when the daily observations were made and bars indicate value range in the 12-hour intervals. Each modeled value shown for (**c-f**; stomatal conductance [mol H_2_O m^-2^s^-1^], **(b)** leaf-level photosynthetic rate [umol CO_2_ m^-2^ s^-1^], **(c)** leaf water potential [Mpa], and **(d)** transpiration [mmol m^-2^ s^-1^]) is averaged for the time interval corresponding to when gas exchange and leaf water potential measurements took place and bars indicate modeled value range. Circle: WW (well-watered); triangle: D1 (drought scenario 1); square: D2 (drought scenario 2).

**Figure S13.**
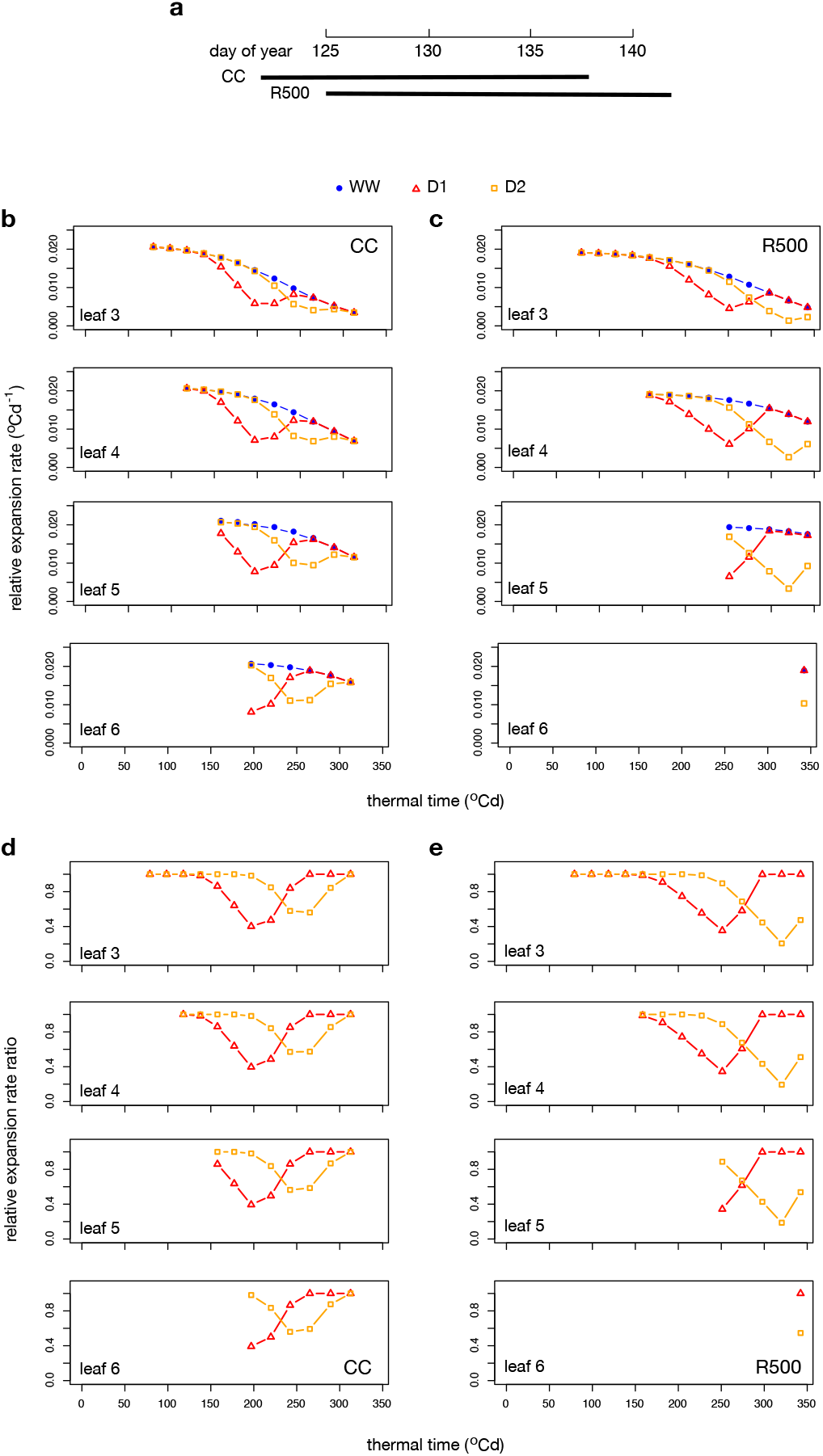
Modulation of relative rate of leaf expansion in response to water limitation. Simulation results from model runs for CC and R500 under WW, D1, and D2 conditions. Plot **(a)** shows the time periods in Julian Day that correspond to the thermal times in **(b-e). (b-c)** Local relative rate of leaf expansion (RER) (^o^Cd^-1^) and **(d-e)** the ratio between RER under the water-limited scenario to RER under the well-watered scenario. X-axes in **(b-e)** show thermal time from the onset of simulation when leaf three initiates.

**Figure S14.**
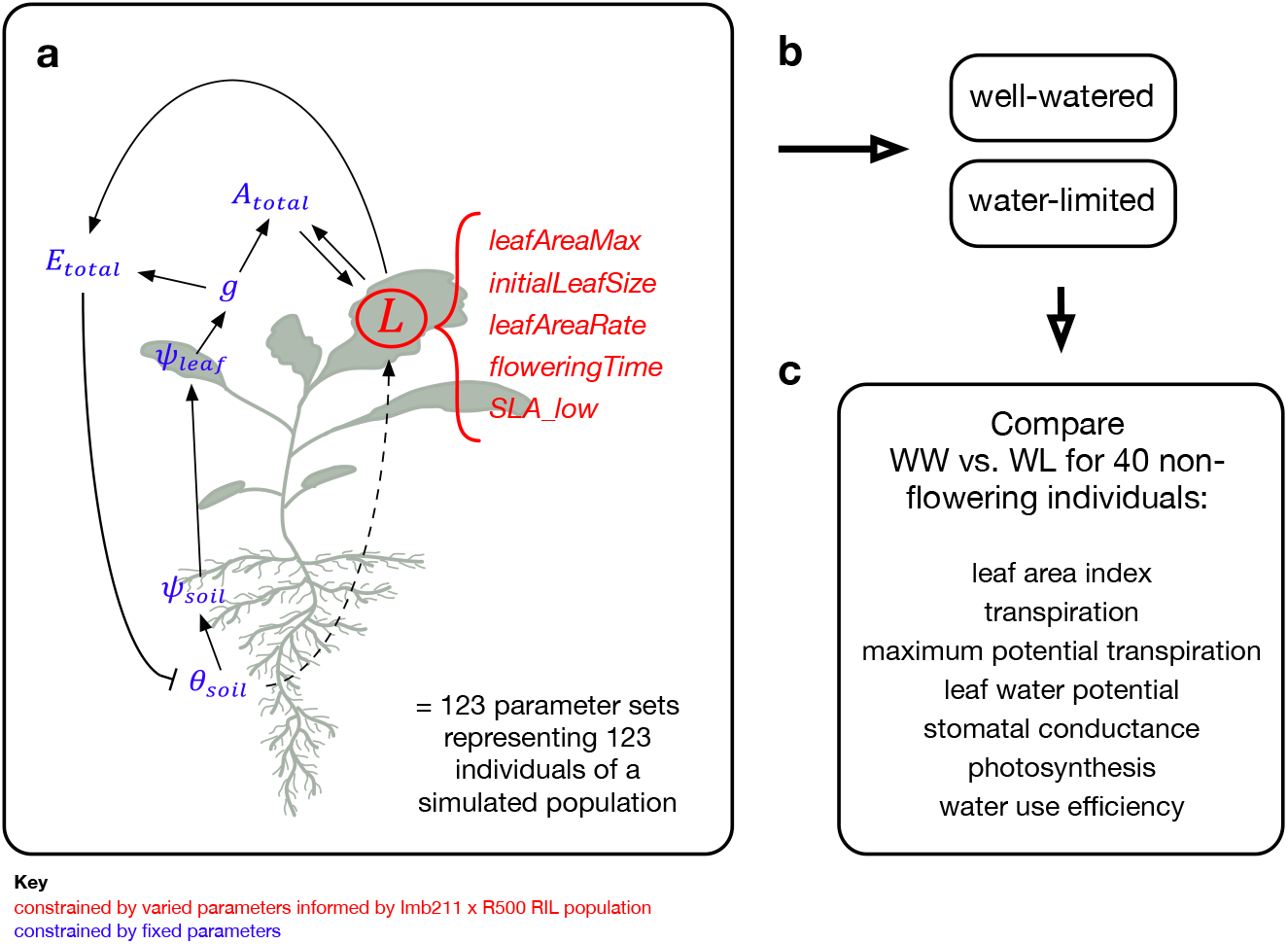
Model exploration using a simulated population. **(a)** Construction of an *in silico* population of 123 individuals using a combination of fixed and varied parameters. Five leaf growth module parameters were varied using values from a real Recombinant Inbred Line population (Imb211 x R500) while other parameters were held constant across the individuals. **(b)** The population was simulated under two scenarios, well-watered (WW) and water-limited (WL), that differed only in water input (see **Figure S15** for environmental drivers). **(c)** 40 simulated individuals that remained in vegetative phase during the simulation were further analyzed.

**Figure S15.**
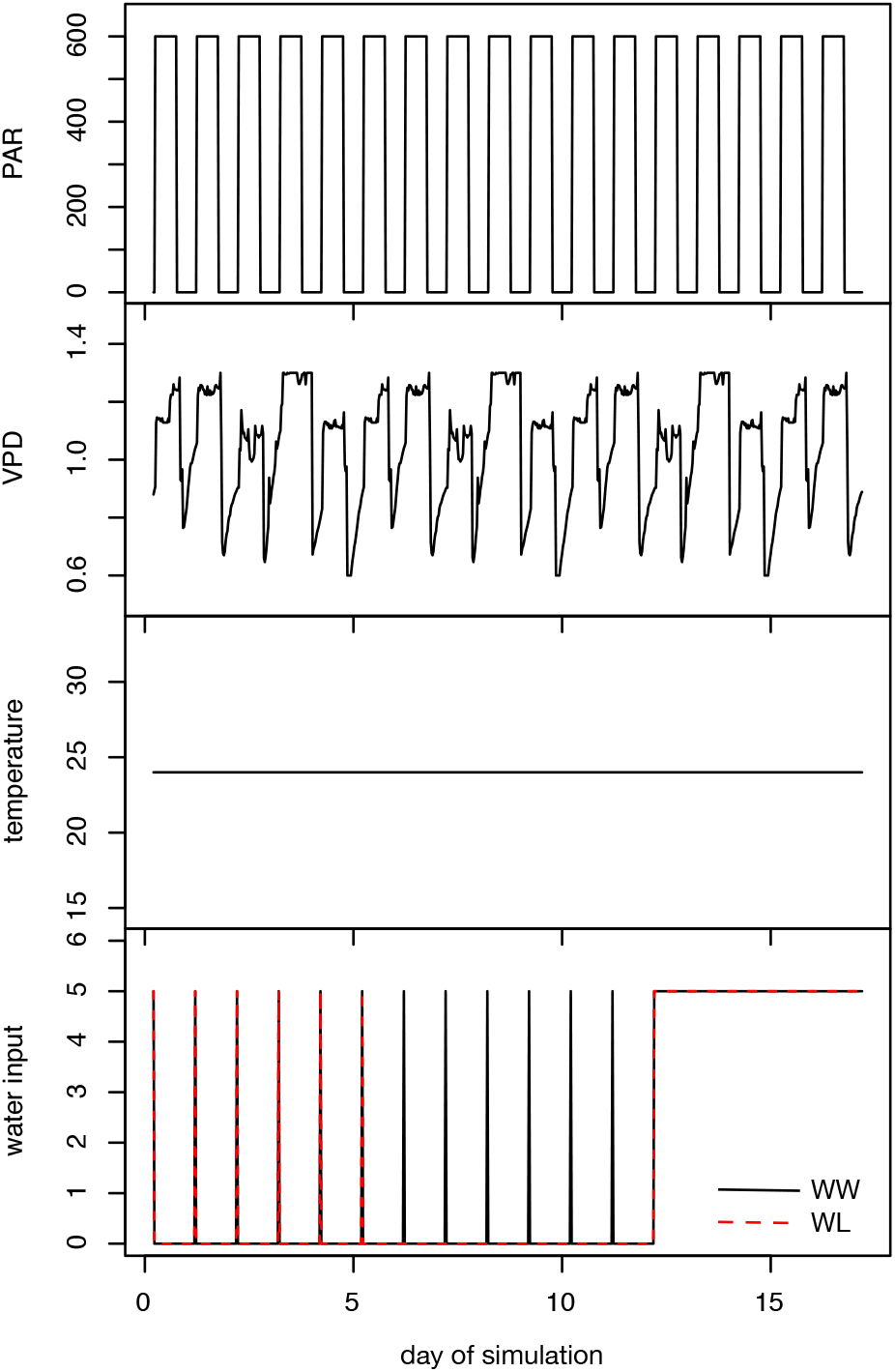
Environmental drivers used for modeling population responses to water deficit. Photosynthetically active radiation (PAR; umol photon m^-2^ s^-1^), vapor pressure deficit (VPD kPa), temperature (^o^C), and water input (mm) shown here.

**Figure S16.**
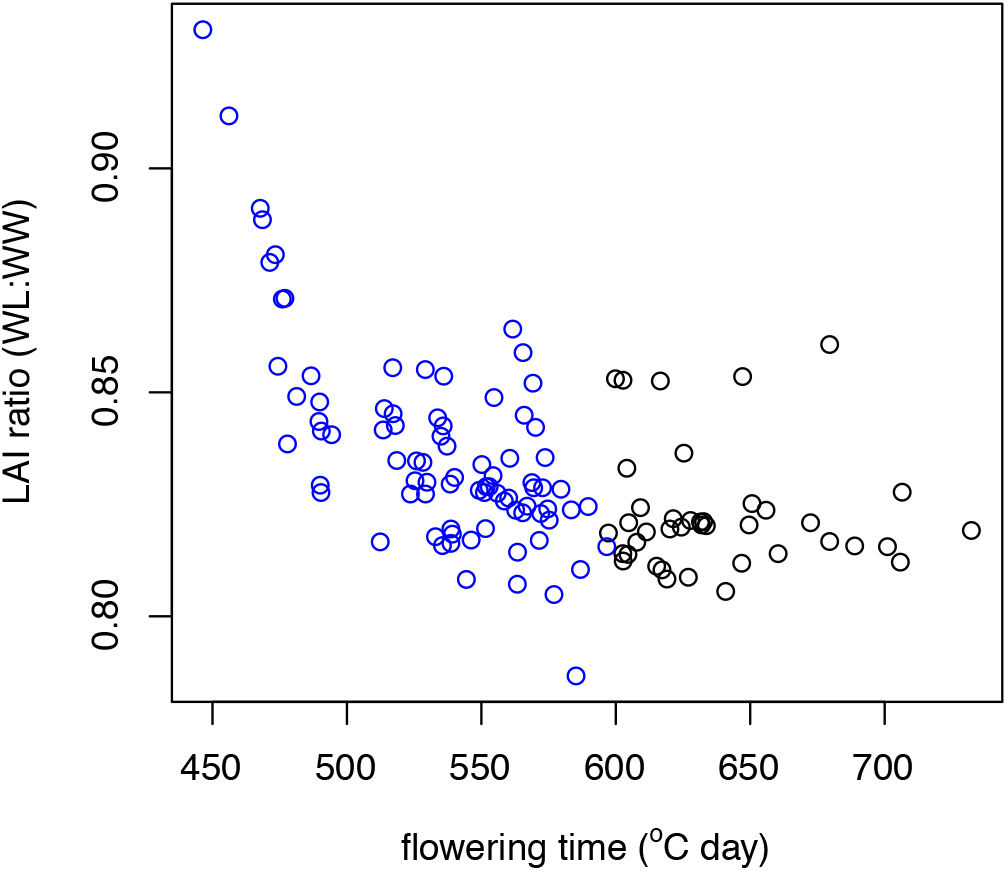
Relationship of final leaf area index ratio (water-limited to well-watered) and flowering time in simulated population. Negative association observed for genotypes that had reached reproductive transition by the end of the simulation period (blue circles).

**Figure S17.**
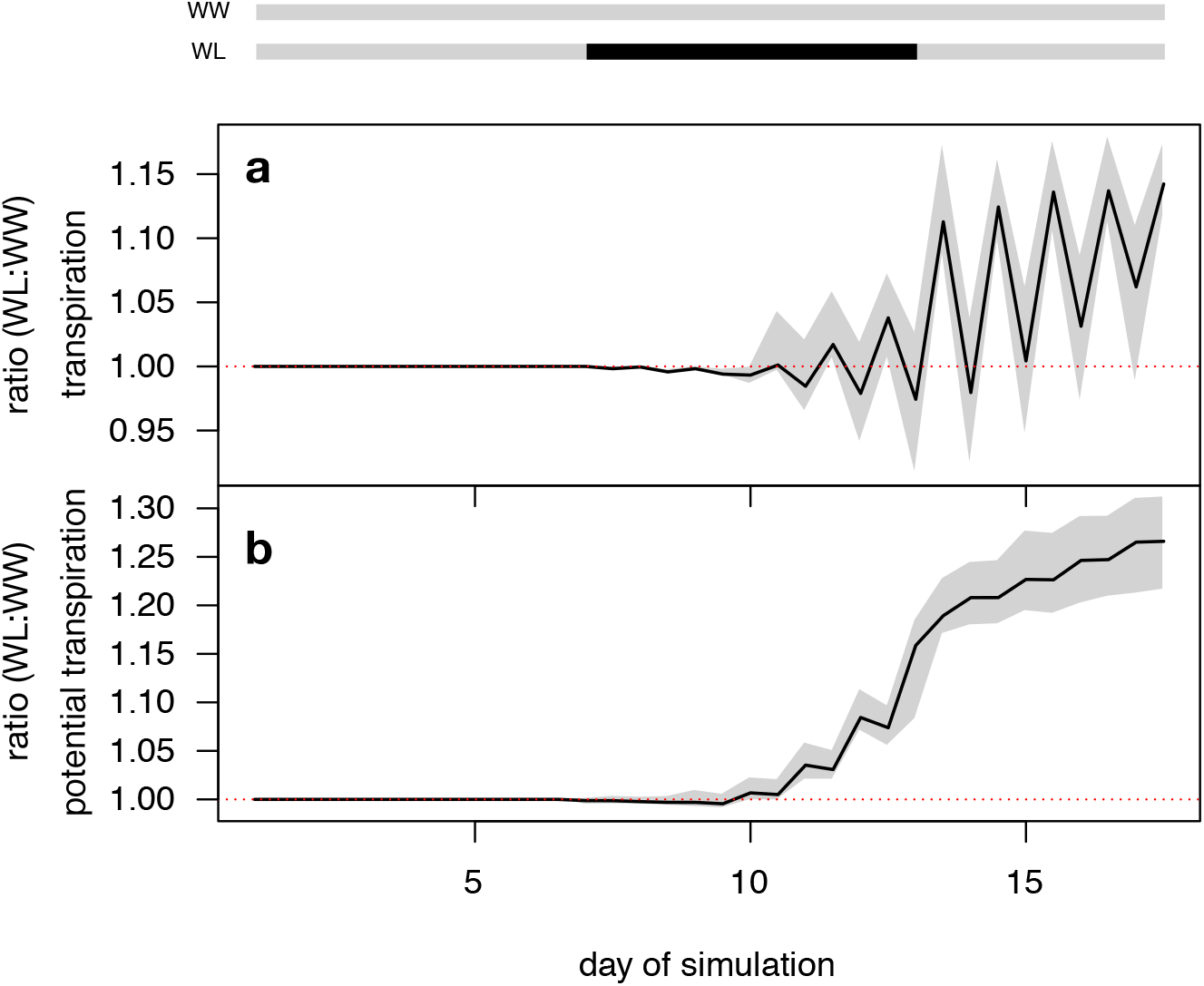
Transpiration of a simulated *B. rapa* population under well-watered and water-limited scenarios. 40 individuals of the *in silico* population that remained in vegetative phase for the entirety of the simulation period are compared here for their behavior under well-watered (WW) and water-limited (WL) conditions. Watering regimes for each scenario are shown as gray and black bars at the top of the figure (gray: daily water input; black: no water input). Pre-dawn (5:00AM timestep) and midday (1:00PM timestep) ratios of WW to WL for **(a)** transpiration (mmol H_2_O m^-2^ s^-1^) and **(b)** modeled maximum potential transpiration (mmol H_2_O m^-2^ s^-1^). Environmental drivers for these simulations are found in **Figure S15**. Shaded gray region: range of values across the 40 individuals; solid black curve: mean. Dotted line: 1.0, where WW and WL have the same value.

**Figure S18.**
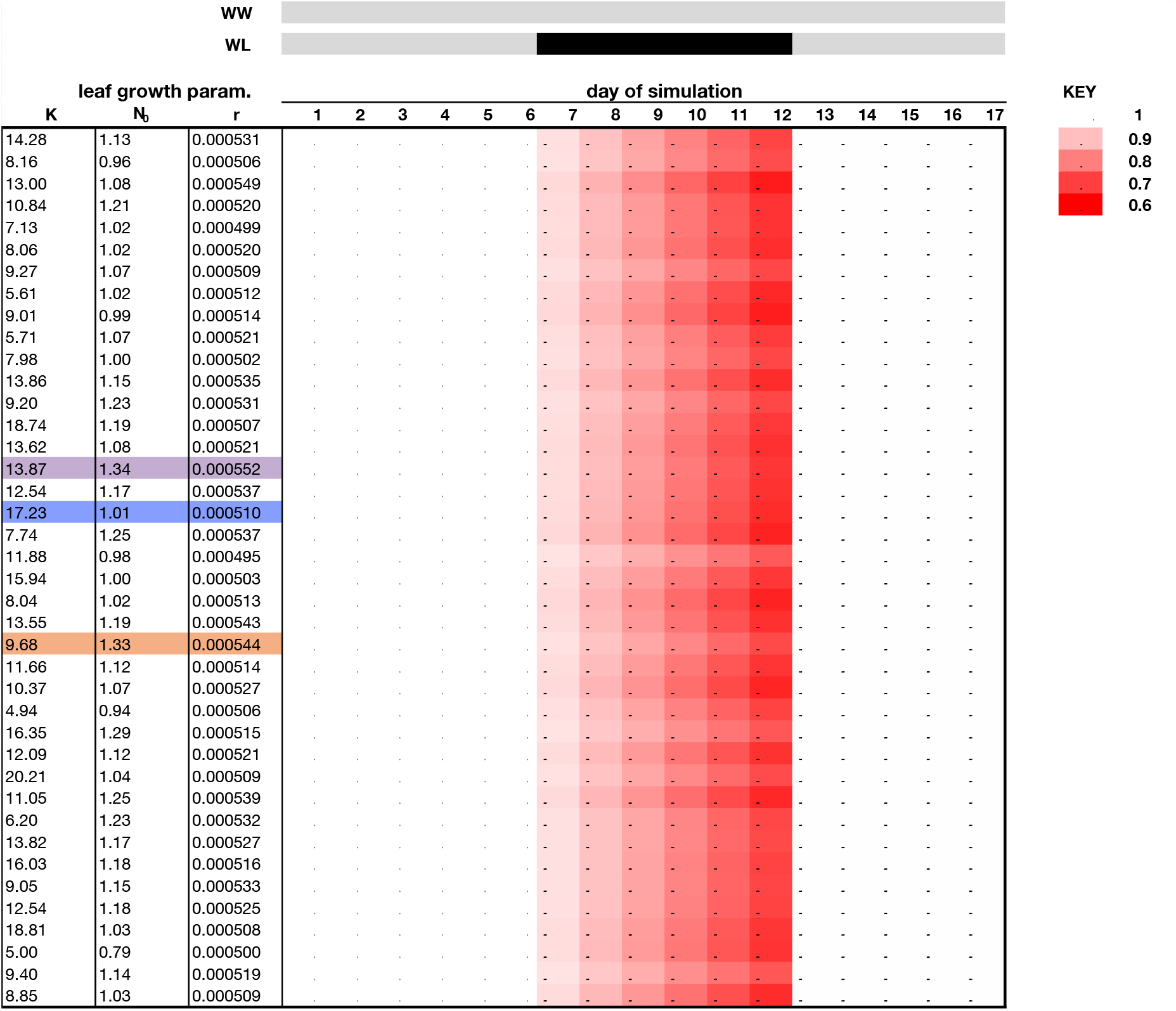
Heat map of volumetric soil water content ratios (WL:WW) across simulation time. Rows show the 40 individuals of the *in silico* population that are examined in **Figure 5**. Colors represents magnitude and direction of water-limited (WL) to well-watered (WW) ratios of modeled volumetric soil water content (%), where red colors indicate lower values in the WL group while blue value indicate higher values in the WL group as compared to its WW counterpart (see Key). Purple, blue, and orange rows mark the individual time-series shown in **Figure 5** with these same colors.

**Figure S19.**
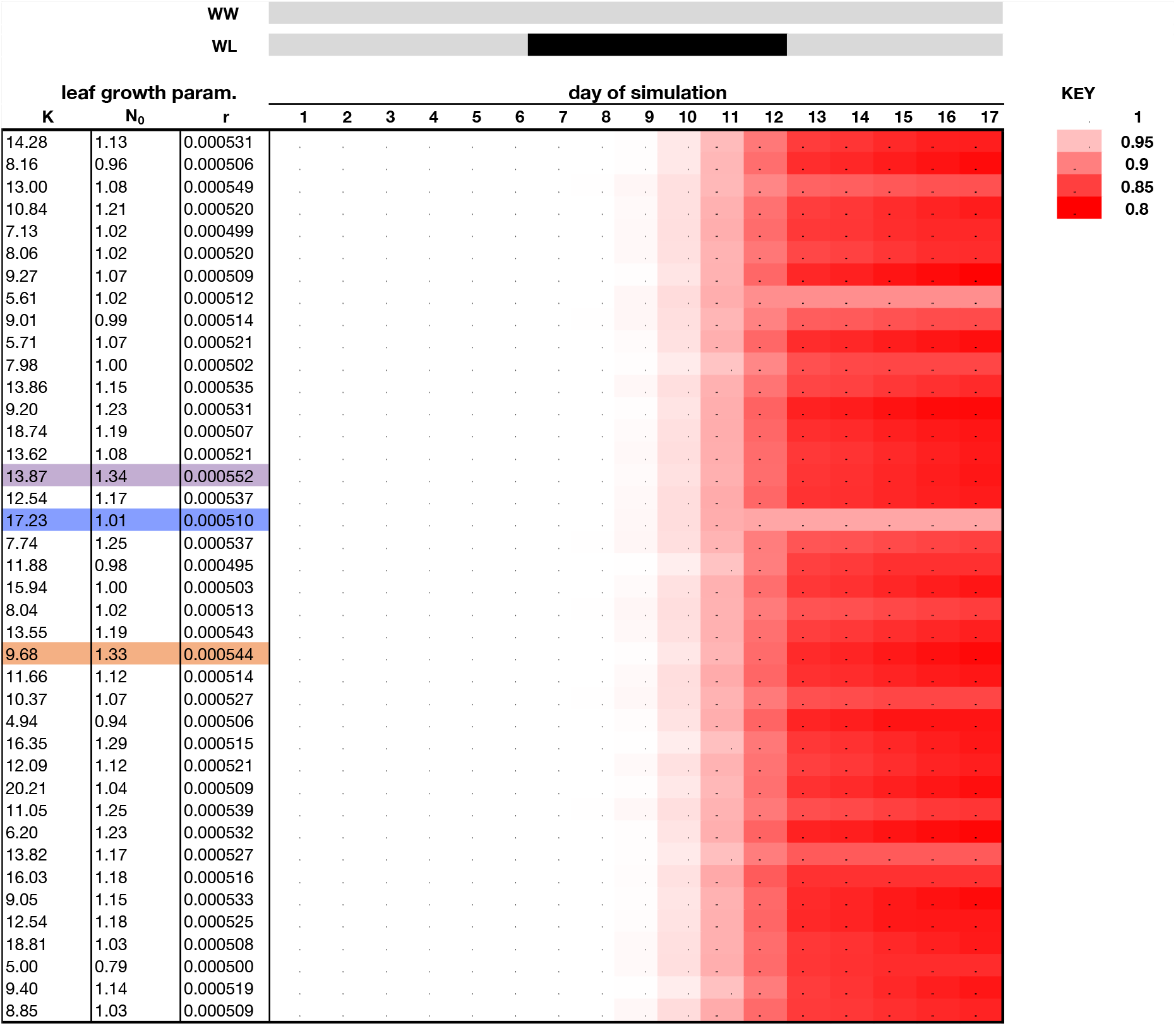
Heat map of leaf area index ratios (WL:WW) across simulation time. Rows show the 40 individuals of the *in silico* population that are examined in **Figure 5**. Colors represents magnitude and direction of water-limited (WL) to well-watered (WW) ratios of modeled leaf area index (cm^2^ cm^-2^), where red colors indicate lower values in the WL group while blue value indicate higher values in the WL group as compared to its WW counterpart (see Key). Purple, blue, and orange rows mark the individual time-series shown in **Figure 5** with these same colors.

**Figure S20.**
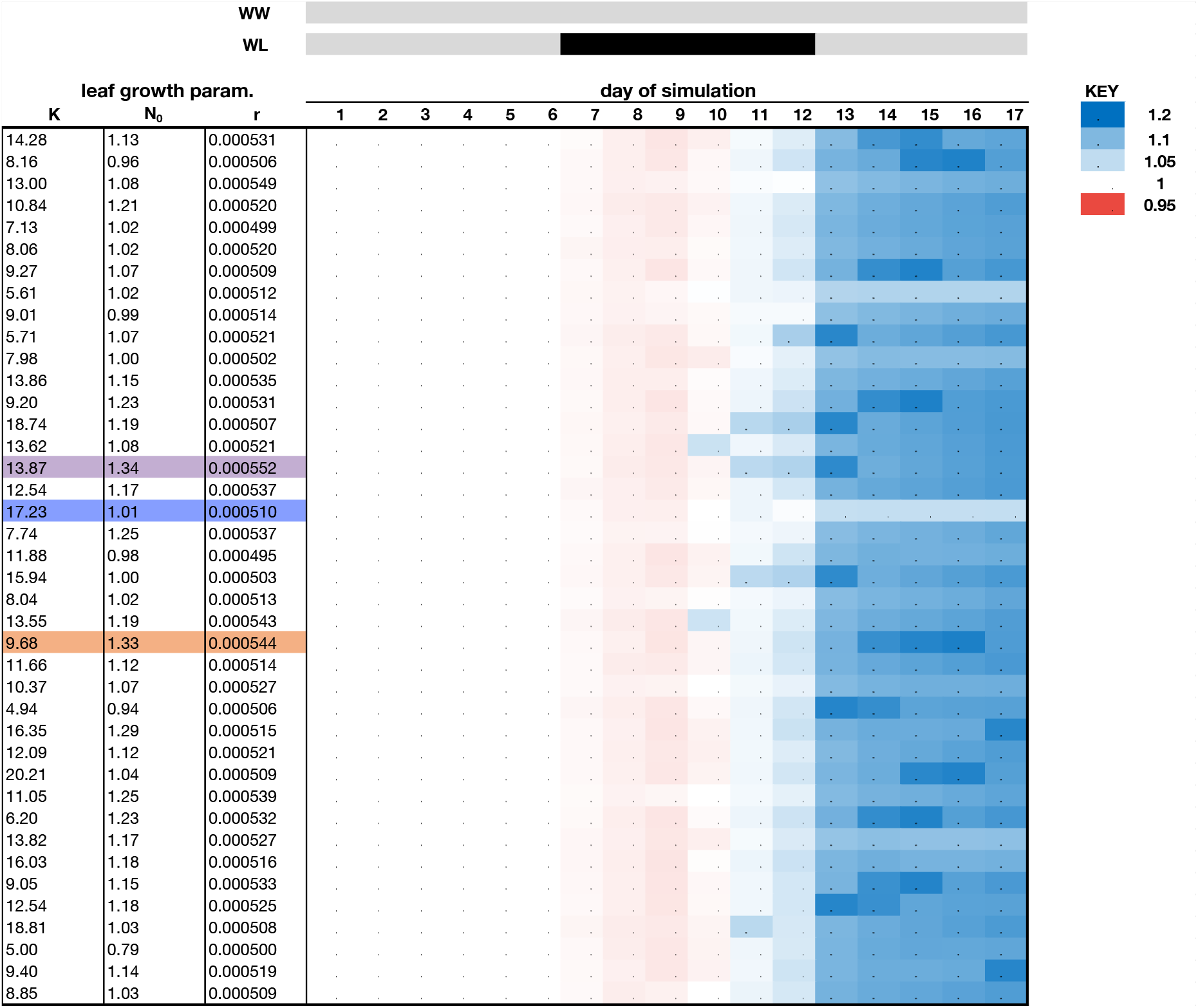
Heat map of midday transpiration ratios (WL:WW) across simulation time. Rows show the 40 individuals of the *in silico* population that are examined in **Figure 5**. Colors represents magnitude and direction of water-limited (WL) to well-watered (WW) ratios of modeled canopy transpiration (mmol H_2_O m^-2^ s^-1^), where red colors indicate lower values in the WL group while blue value indicate higher values in the WL group as compared to its WW counterpart (see Key). Purple, blue, and orange rows mark the individual time-series shown in **Figure 5** with these same colors.

**Figure S21.**
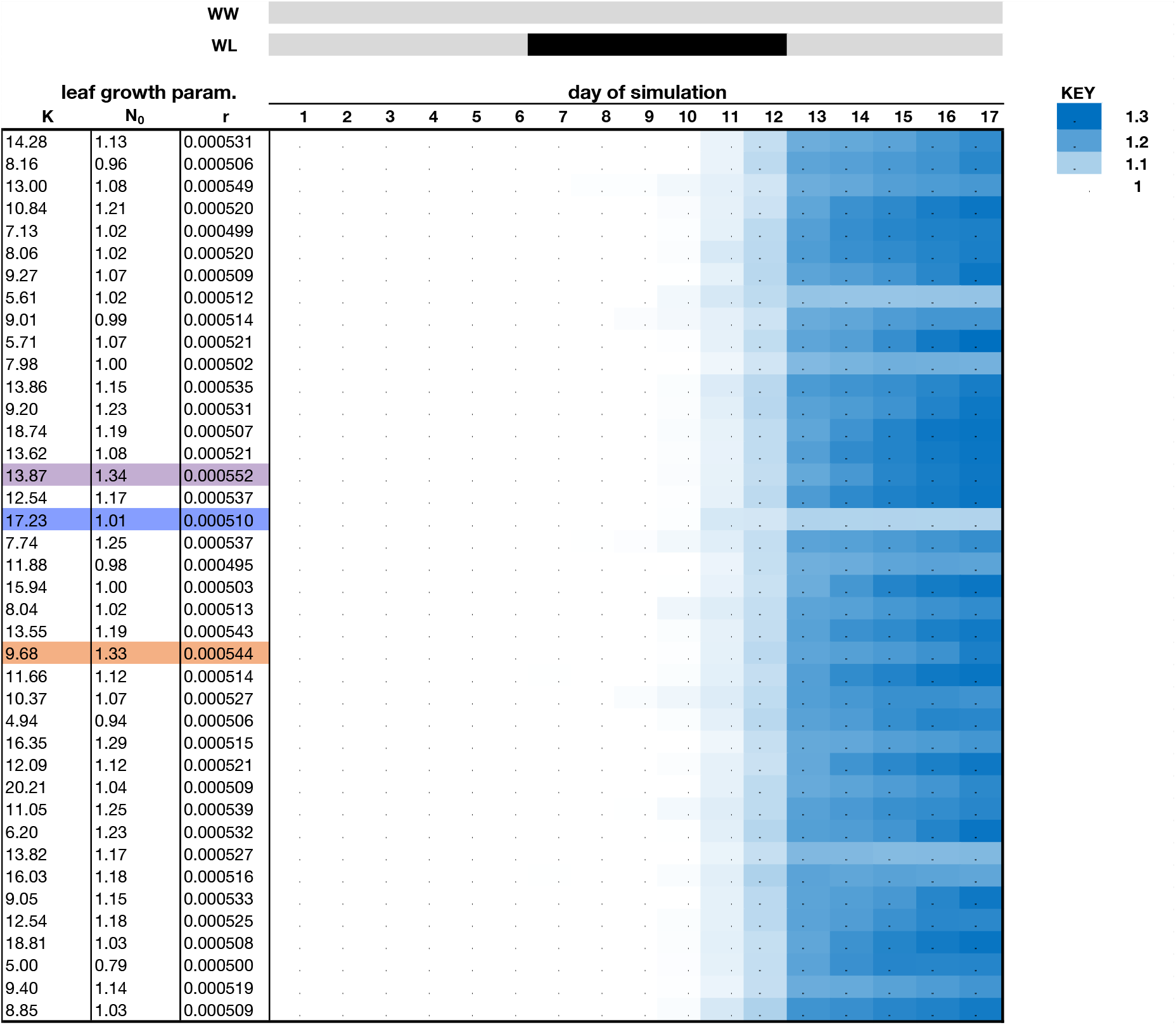
Heat map of midday maximum transpiration ratios (WL:WW) across simulation time. Rows show the 40 individuals of the *in silico* population that are examined in **Figure 5**. Colors represents magnitude and direction of water-limited (WL) to well-watered (WW) ratios of modeled canopy maximum transpiration (mmol H_2_O m^-2^ s^-1^), where red colors indicate lower values in the WL group while blue value indicate higher values in the WL group as compared to its WW counterpart (see Key). Purple, blue, and orange rows mark the individual time-series shown in **Figure 5** with these same colors.

**Figure S22.**
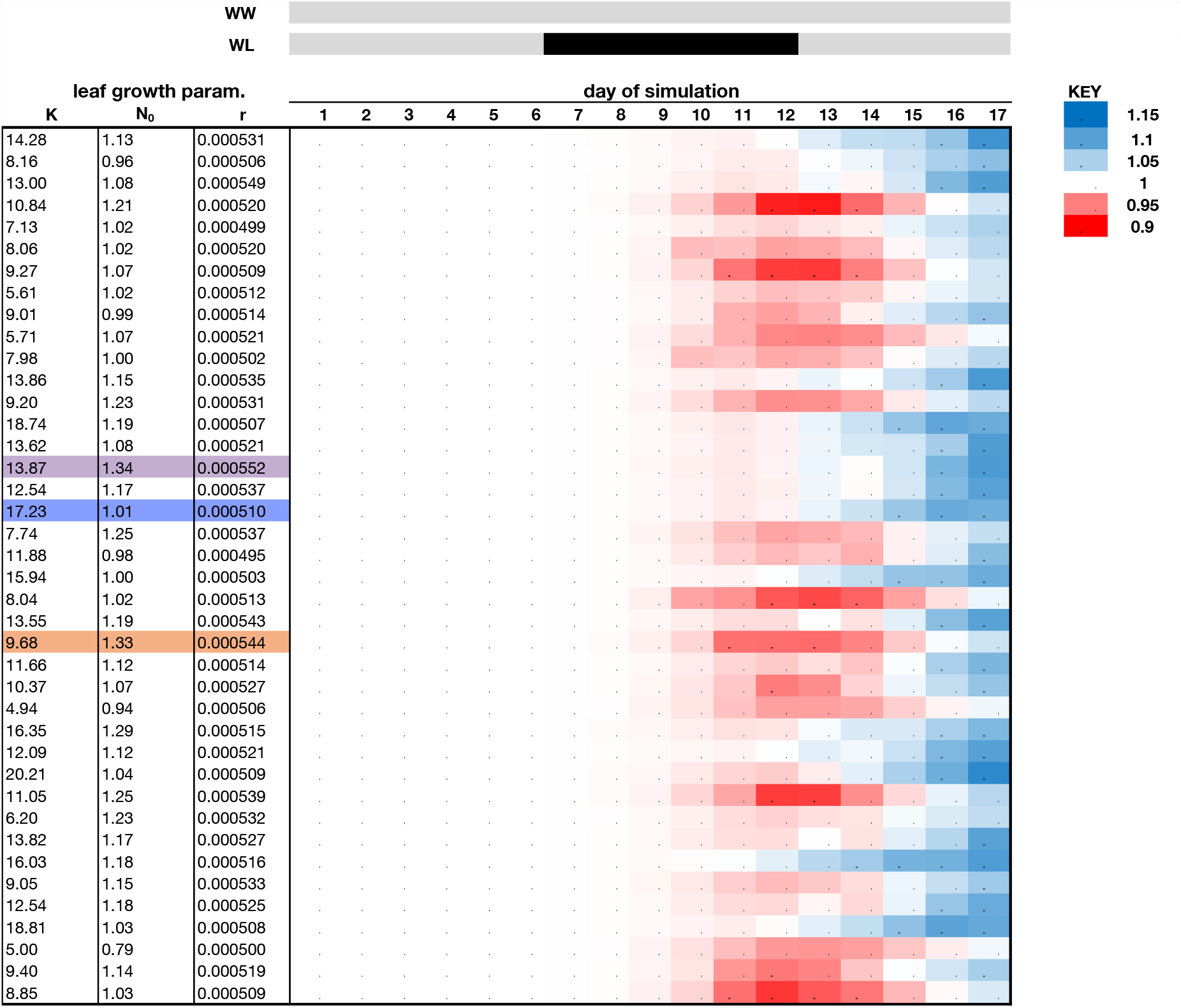
Heat map of midday photosynthesis ratios (WL:WW) across simulation time. Rows show the 40 individuals of the *in silico* population that are examined in **Figure 5**. Colors represents magnitude and direction of water-limited (WL) to well-watered (WW) ratios of modeled canopy photosynthesis (umol CO_2_ m^-2^ s^-1^), where red colors indicate lower values in the WL group while blue value indicate higher values in the WL group as compared to its WW counterpart (see Key). Purple, blue, and orange rows mark the individual time-series shown in **Figure 5** with these same colors.

**Figure S23.**
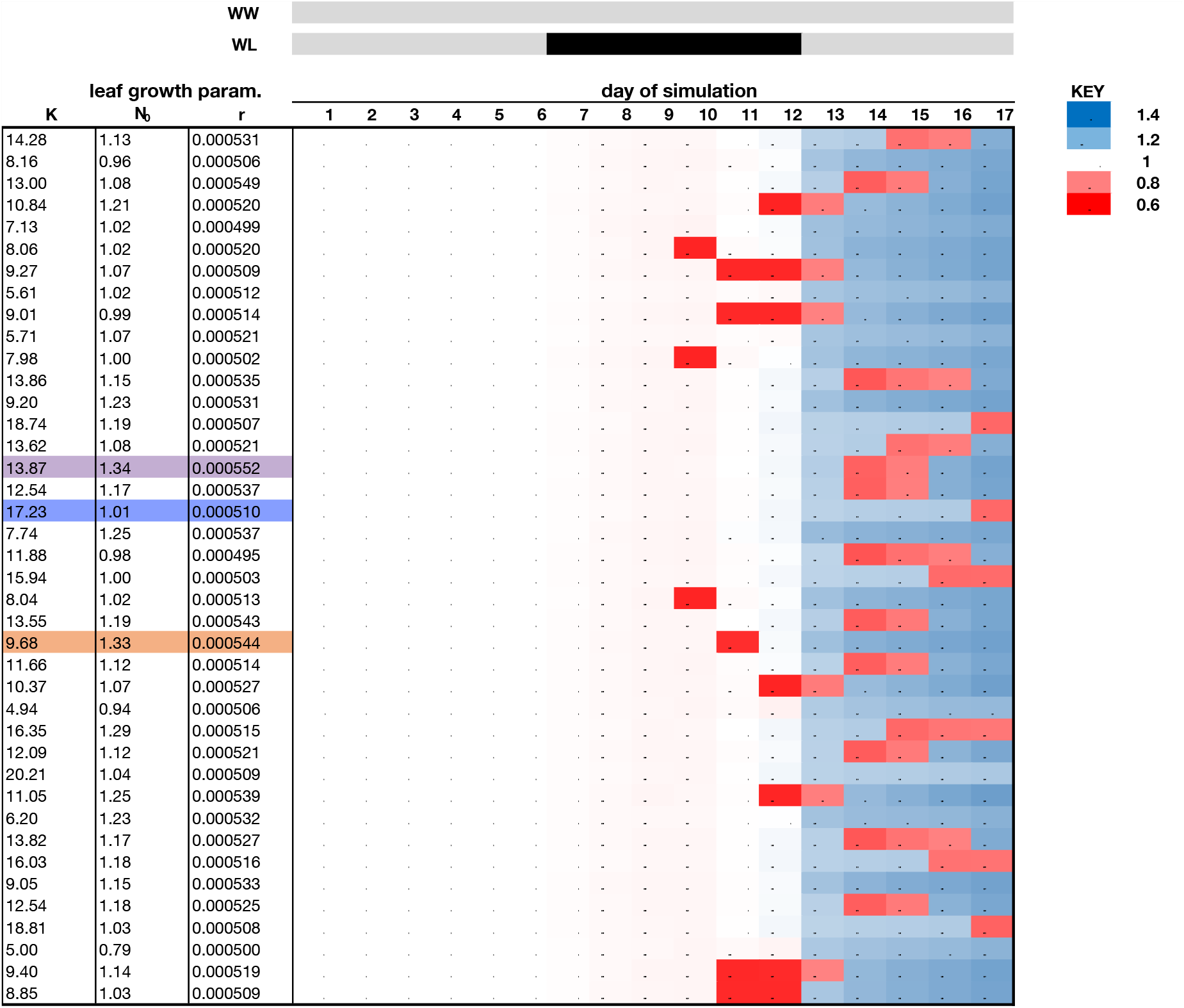
Heat map of midday stomatal conductance ratios (WL:WW) across simulation time. Rows show the 40 individuals of the *in silico* population that are examined in **Figure 5**. Colors represents magnitude and direction of water-limited (WL) to well-watered (WW) ratios of modeled canopy stomatal conductance (mol H_2_O m^-2^ s^-1^), where red colors indicate lower values in the WL group while blue value indicate higher values in the WL group as compared to its WW counterpart (see Key). Purple, blue, and orange rows mark the individual time-series shown in **Figure 5** with these same colors.

**Figure S24.**
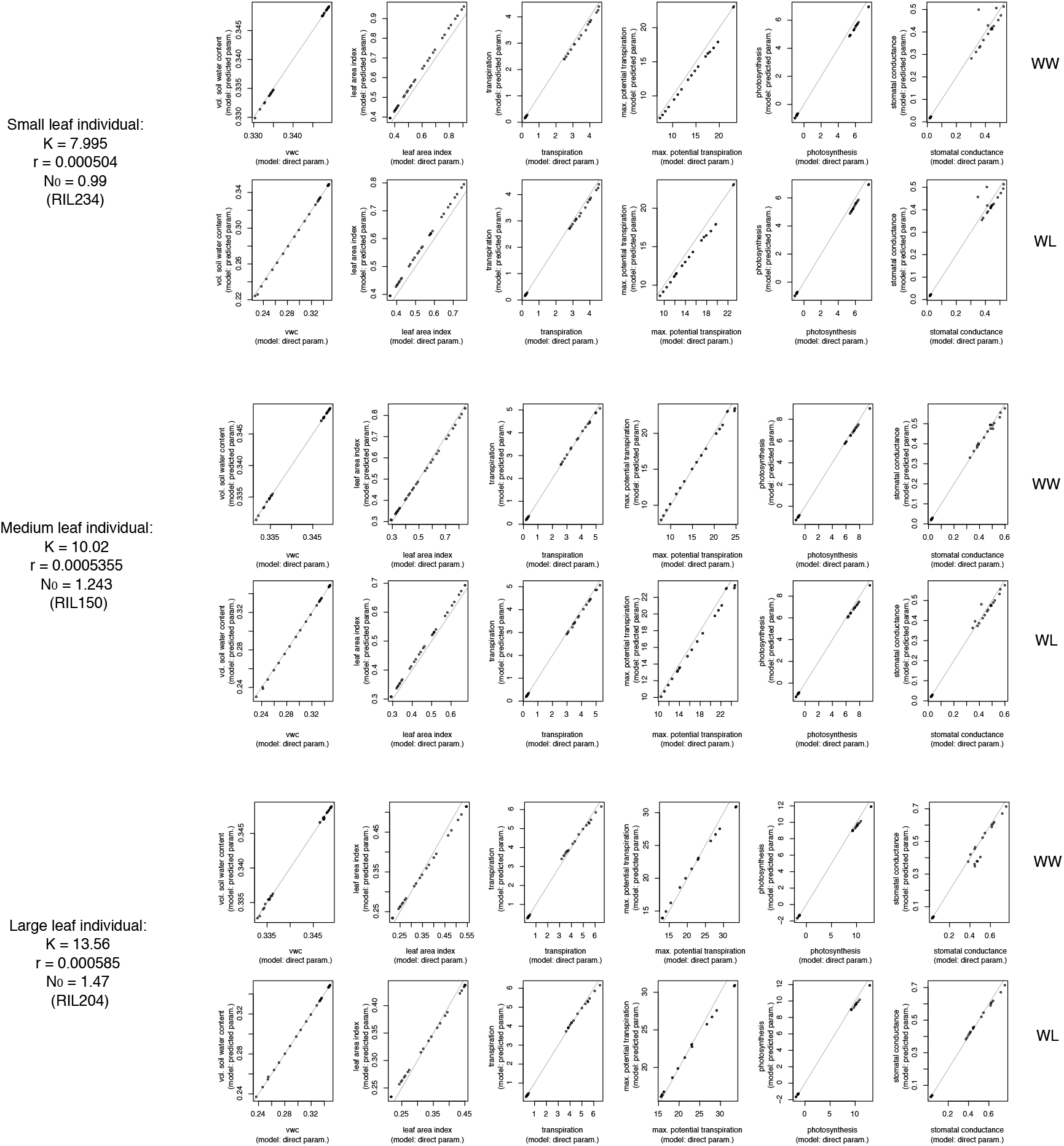
Comparison of simulation results informed by directly estimated parameters versus informed by predicted parameters. From the *in silico* population of 123 individuals, three were selected based on maximum leaf area (chosen at 25, 50, and 75% quantiles). Stochastic simulations were performed (25 simulations x 3 individual x 2 treatments x method (predicted versus direct): 300 simulations). Traits examined are predawn (5:00AM timestep) and midday (1:00PM timestep) values of volumetric soil water content (%), leaf area index (cm^2^ cm^-2^), canopy transpiration (mmol H_2_O m^-2^ s^-1^), canopy maximum potential transpiration (mmol H_2_O m^-2^ s^-1^), canopy photosynthesis (umol CO_2_ m^-2^ s^-1^), and canopy stomatal conductance (mol H_2_O m^-2^ s^-1^).

**Table S1.** Summary table expansion of Figure 1a.

| Stage | Description | Plant material | Data/information collected | Source | Cross-reference to figures and tables |
| --- | --- | --- | --- | --- | --- |
| 1 | model development | -- | theoretical growth equation | literature | Figure 1b, Table S2-S3, Appendix S1 |
| 2a | observations under non-stressed conditions | *B. rapa* morphotypes: vegetable turnip (VT), chinese cabbage (CC) and oilseed (R500) | leaf growth dynamics | this study | Figure S1, S3, |
| 2b | genomic prediction of input parameters | BraIRRI population | leaf growth dynamics (on 2nd leaf) and SNP data | literature | Figure 2, Figure S8, Table S4 |
| 3 | model update | -- | -- | -- | Figure S4-S7 |
| 4 | evaluation under mild stress conditions | genotypes CC and R500 | leaf growth dynamics and suite of physiological traits | this study | Figure 3, Figure S2, S9-S10, Table S5 |
| 5 | model update | -- | -- | -- | Figure 4, Figure S11-S13 |
| 6 | model exploration on a simulated population | a simulated population informed by real population leaf growth parameters | -- | -- | Figure 5, Figure S14-S24 |

**Table S2.** Growth parameter values of *B. rapa* morphotypes used for TREES modeling.

| Parameter | R500 | VT | CC | Evaluation method |
| --- | --- | --- | --- | --- |
| leafAreaMax | 19.65 | 29.88 | 38.59 | experimental data |
| initialLeafSize | 0.25 | 0.4 | 0.65 | experimental data |
| leafArea_Rate | 0.0008132 | 0.0008904 | 0.0008926 | experimental data |
| Kalpha | 334.13 | 694.72 | 883.31 | experimental data |
| Kbeta | 17.00 | 23.22 | 22.87 | experimental data |
| Nalpha | 4.45 | 4.50 | 8.11 | experimental data |
| Nbeta | 16.52 | 10.47 | 11.89 | experimental data |
| ralpha | 68.70 | 74.81 | 118.31 | experimental data |
| rbeta | 83996.62 | 83027.69 | 131431.51 | experimental data |
| dur_LeafExpansion | 11024.02 | 9985.409 | 9708.083 | experimental data |
| SLA_max, SLA_low | 14, 2142 | 14, 2142 | 14, 2142 | estimated <sup>1</sup> |
| leaf_insertAngle | 60 | 45 | 85 | experimental data |
| leaf_len_to_width | 2.5 | 2 | 1.25 | experimental data |
| phyllochron | 1970.534 | 1979.803 | 1026.731 | experimental data |
| floweringTime | 23850 | 50000 | 50000 | estimated <sup>2</sup> |
| Tbase | 0.96 | 0.96 | 0.96 | literature value |
| therm_plant_init | 4926.335 | 4949.5075 | 2566.8275 | set to 2.5 times phyllochron |
| projectedArea_init | 15 | 15 | 15 | estimated <sup>3</sup> |
| pot_size | 40 | 40 | 40 | experimental data |
| root_to_shoot | 0.43 | 0.26 | 0.24 | experimental data |
| leaf_to_stem | 1.59 | 15.4 | 35.4 | experimental data |

Note: Units for parameters found in Table 1. ‘alpha’ and ‘beta’ parameters refer to gamma distribution rate and shape parameters, respectively.
<sup>1</sup>no experimental data collected; SLA\_low from literature; SLA\_max (only used in initialization along with projectedArea\_init) was tuned
<sup>2</sup>R500 measured directly; VT and CC did not flower prior to end of experiment, so a large number was selected for simulations
<sup>3</sup>no experimental data collected so all genotypes were set to same value; requires some tuning with SLA\_max

**Table S3.** Maximum leaf sizes across leaf number.

**a**
| leaf | CC |  |  | VT |  |  | R500 |  |  |
| --- | --- | --- | --- | --- | --- | --- | --- | --- | --- |
|  | mean | se | N | mean | se | N | mean | se | N |
| 1 | 16.44 | 3.37 | 5 | 7.58 | 2.08 | 5 | 5.48 | 0.44 | 16 |
| 2 | 12.90 | 2.50 | 5 | 5.02 | 0.58 | 5 | 7.72 | 0.45 | 16 |
| 3 | 34.93 | 4.14 | 5 | 20.10 | 2.63 | 5 | 14.18 | 0.93 | 16 |
| 4 | 48.87 | 2.23 | 5 | 25.33 | 5.44 | 5 | 20.56 | 1.39 | 16 |
| 5 | 51.88 | 3.20 | 5 | 43.69 | 6.41 | 5 | 18.76 | 1.94 | 16 |
| 6 | 54.31 | 1.61 | 5 | 47.61 | 8.40 | 5 | 17.94 | 3.19 | 16 |
| 7 | 47.34 | 4.85 | 5 | 34.74 | 8.44 | 5 | -- | -- | -- |
| 8 | 45.55 | 3.81 | 5 | 28.90 | 14.64 | 5 | -- | -- | -- |

**b**
| leaf | RIL301 |  |  | RIL46 |  |  |
| --- | --- | --- | --- | --- | --- | --- |
|  | mean | se | N | mean | se | N |
| 1 | 4.25 | 0.19 | 5 | 1.82 | 0.15 | 5 |
| 2 | 6.44 | 0.42 | 5 | 2.32 | 0.45 | 5 |
| 3 | 10.53 | 0.76 | 5 | 2.93 | 0.61 | 5 |
| 4 | 13.12 | 0.93 | 5 | 3.10 | 0.52 | 5 |
| 5 | 8.27 | 1.24 | 5 | 2.61 | 0.25 | 5 |

'se': standard error of the mean
'N': number of individual plants surveyed

**Table S4.**
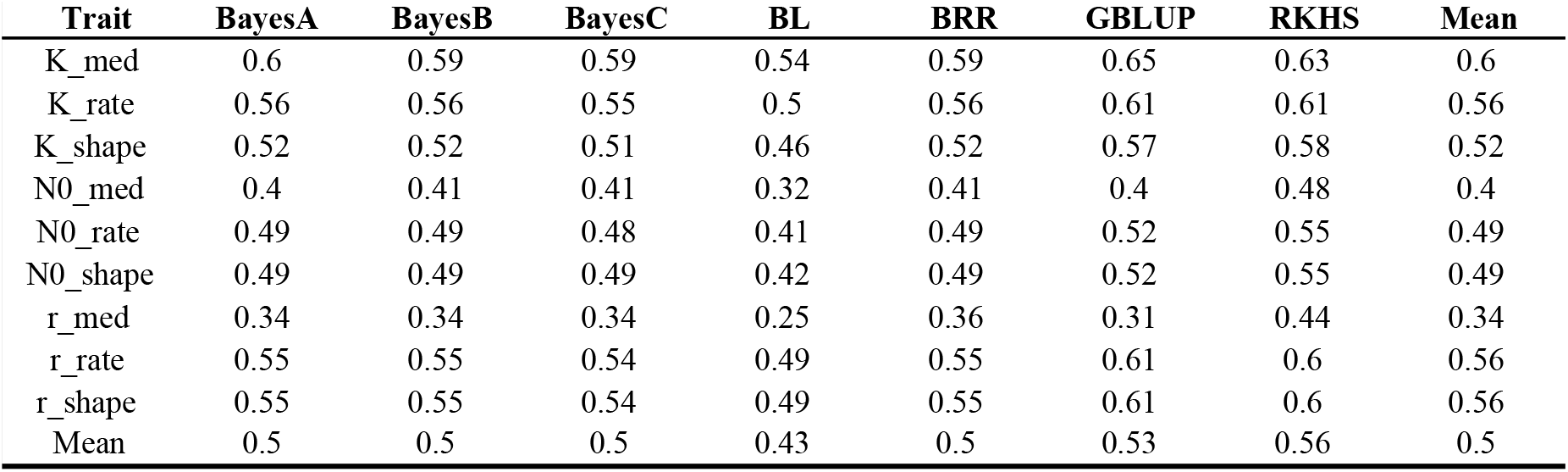
Genomic heritability of leaf growth parameters.

| **Trait** | **BayesA** | **BayesB** | **BayesC** | **BL** | **BRR** | **GBLUP** | **RKHS** | **Mean** |
| --- | --- | --- | --- | --- | --- | --- | --- | --- |
| K_med | 0.6 | 0.59 | 0.59 | 0.54 | 0.59 | 0.65 | 0.63 | 0.6 |
| K_rate | 0.56 | 0.56 | 0.55 | 0.5 | 0.56 | 0.61 | 0.61 | 0.56 |
| K_shape | 0.52 | 0.52 | 0.51 | 0.46 | 0.52 | 0.57 | 0.58 | 0.52 |
| N0_med | 0.4 | 0.41 | 0.41 | 0.32 | 0.41 | 0.4 | 0.48 | 0.4 |
| N0_rate | 0.49 | 0.49 | 0.48 | 0.41 | 0.49 | 0.52 | 0.55 | 0.49 |
| N0_shape | 0.49 | 0.49 | 0.49 | 0.42 | 0.49 | 0.52 | 0.55 | 0.49 |
| r_med | 0.34 | 0.34 | 0.34 | 0.25 | 0.36 | 0.31 | 0.44 | 0.34 |
| r_rate | 0.55 | 0.55 | 0.54 | 0.49 | 0.55 | 0.61 | 0.6 | 0.56 |
| r_shape | 0.55 | 0.55 | 0.54 | 0.49 | 0.55 | 0.61 | 0.6 | 0.56 |
| Mean | 0.5 | 0.5 | 0.5 | 0.43 | 0.5 | 0.53 | 0.56 | 0.5 |

**Table S5.** Observed ratio of shoot to root under mild drought.

| **Genotype** | **Treatment** | **mean** | **se** | **N** |
| --- | --- | --- | --- | --- |
| **CC** | **D1** | 13.199 | 1.453 | 47 |
|  | **D2** | 11.004 | 0.696 | 52 |
|  | **WW** | 17.096 | 2.506 | 26 |
| **R500** | **D1** | 5.969 | 0.364 | 48 |
|  | **D2** | 5.065 | 0.277 | 50 |
|  | **WW** | 9.078 | 0.948 | 25 |

Ratio measured as dry weight of above-ground biomass to belowground biomass. The number of replicates per genotype by treatment combination found in the column labeled 'N'; se: standard error of the mean.

## Appendix S1. TREES leaf growth sub-model appendix

The new canopy sub-model in Terrestrial Regional Ecosystem Exchange Simulator (TREES)^1^ developed in this study adds 22 additional input parameters (**Table 1**). An overview of the algorithm performed at each 30-minute timestep in simulations is presented below.

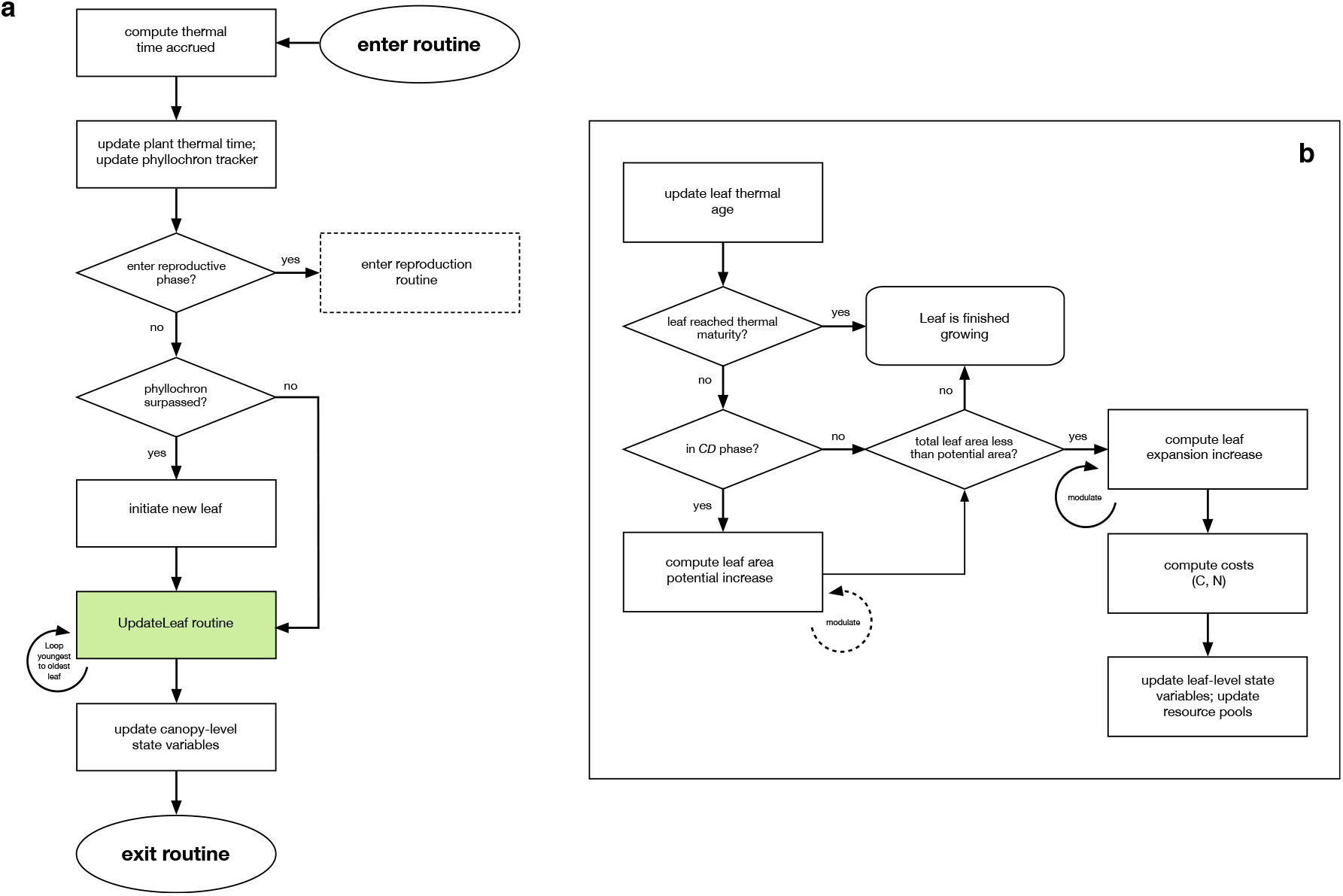
Schematic diagram of the new leaf-based canopy development sub-model in TREES. **(a)** Summary of the overall algorithm for sequential leaf growth and development found in simulation_functions.cpp. **(b)** Steps in the UpdateLeaf function (highlighted as green box in **(a)**), part of the bgc class (bgc.cpp), used to update each emerged leaf per simulated 30 minute time-step. Dashed elements indicate parts of the model that currently contain placeholders as targets for future development.

### Model initialization

At initialization, new model parameters link directly to original TREES state variables, which track within-plant resources such as structural and non-structural carbohydrates (kg C/ha) and nitrogen (kg N/ha) over time. The parameter total projected shoot area is divided by the parameter, *SLA_max_* (m^2^/kg C), to provide a non-destructive estimation of total plant biomass to initialize model resource pools. Projected shoot area at third leaf emergence stage was used for initialization. *SLA_max_* represents the specific leaf area (SLA) of leaves at initiation and likely requires some tuning due to the fact that direct measurements typically destructive. Estimated total aboveground plant biomass is then decomposed into leaf and stem to initialize model resource pools.

### Leaf growth

Thermal-dependent growth of an individual leaf proceeds in two phases partially overlapping in time. The first phase represents cell division (CD), i.e., establishment of leaf area potential. The second phase represents leaf area expansion (EXP), i.e., realization of leaf area potential. We model both CD and EXP with the same governing logistic growth function, parameterized by inputs from direct observation (or genomic prediction) of the EXP phase. This simplification is supported by previous work that documented similar relationships between cell division and expansion to thermal time in sunflower leaves^2^. At each simulated time-step, the theoretical relative rate of growth is computed and consequently scaled by available resources to give rise to a realized growth increment of leaf area potential, actual leaf area, or both. Theoretical relative rate of growth of leaf *i* during phase *j* follows

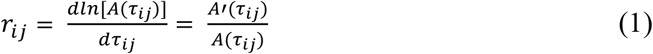

where *τ_ij_* is thermal time of leaf *i* spent in process *j* (either CD or EXP). *A*(*τ_ij_*) is the thermal time-dependent leaf area following a logistic growth function, 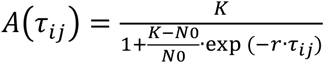, parameterized by inputs *K, N_0_*, and *r*. Because hydraulic conductance measurements in *B. rapa* is challenging to collect at high time resolution, the primary modulation function for leaf area expansion is an empirical relationship based on water availability in the soil and is described

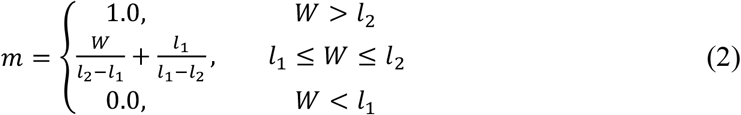

where *m* is a coefficient whose value ranges from 0 to 1. *W* is the soil water content relative to maximum soil water content that is dependent upon soil texture characteristics (original TREES parameters^1^). *l_1_* and *l_2_* delimit a range of *W* within which *c* is scaled linearly to *W*; outside this range, *m* is either 0 or 1. We set *l_1_* and *l_2_* to 0.65 and 0.85, respectively, to approximate observations from the mild drought experiment in CC D2 (Figure S2b). From equations 1 and 2, realized increase in leaf area at each time-step for leaf *i* is

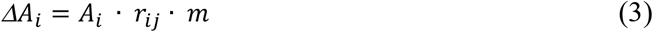

where *j* is EXP phase and *A_i_* is the area of leaf *i*.

### Metabolic costs to leaf growth

For each time-step during which EXP occurs, a specific leaf area for leaf *i, S_i_* (m^2^ kg^-1^), based on plant nitrogen is computed:

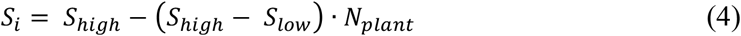

where *S_low_* is an input parameter and *S_high_* is twice that. *N_plant_* ranges from 0 to 1. From equations 3 and 4, structural carbon, AC, required for incremental growth is

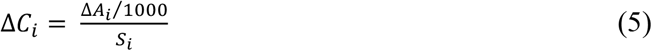

Growth respiration for leaf development is modeled as construction cost (g C substrate g^-1^ C product). Due to impracticalities of future validation for cost to CD, construction cost is taken only in EXP. Construction respiration cost, *R_i_*, is computed

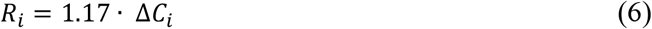

1.17 g C substrate g^-1^ C product for leaf expansion assumes that most of the cost goes to building carbohydrates such as lignin in cell walls^3^. Original TREES routines for maintenance respiration associated with leaf biomass were retained^1^.

### Linkages to canopy-level features, stem and root growth

Leaf-level traits contribute to canopy-level characteristics such as leaf area index and plant-average specific leaf area. For computation simplicity, we assume elliptic shapes for *B. rapa* leaves having a length to width ratio, *c*, and extruding from the main stem axis at an insertion angle, *θ*(^o^). Parameters *c* and 0 may be adjusted to reflect the variation observed across different *B. rapa* genotypes. In the current model, *c* and *θ* remain fixed throughout the simulation of a single genotype but may potentially be adjusted in the future to reflect dynamic leaf shape and angle across leaves or cohorts of leaves within individual plants. Projected area (cm^2^) of a single leaf *i* is computed:

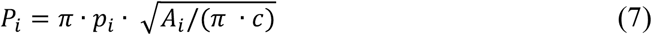

where *A_i_* is the current area of leaf *i* and *p_i_* is the length of the projected ellipse. *p_i_* is computed:

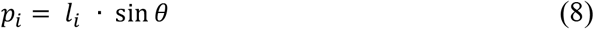

and *l_i_* is computed by assuming an elliptic leaf shape as follows, where *w_i_* is the width of leaf *i*:

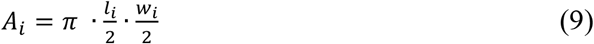

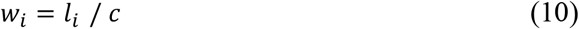

Equations 9 and 10 are combined to solve for *l_i_*:

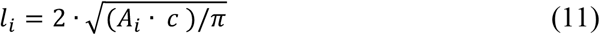

Total projected leaf area (cm^2^) is computed by summing the parameter initial projected shoot area, *A_init_* and the projected leaf area, *P_i_*, across *n* emerged leaves:

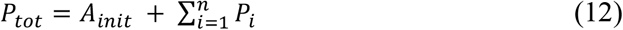

LAI (m^2^ m^-2^), *L*, is computed:

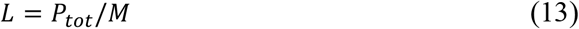

where *M* is the maximum projected ground area (cm^2^), calculated as the area of a circle with radius *p_i_* when *A_i_* is maximum. A graphical explanation is provided below.

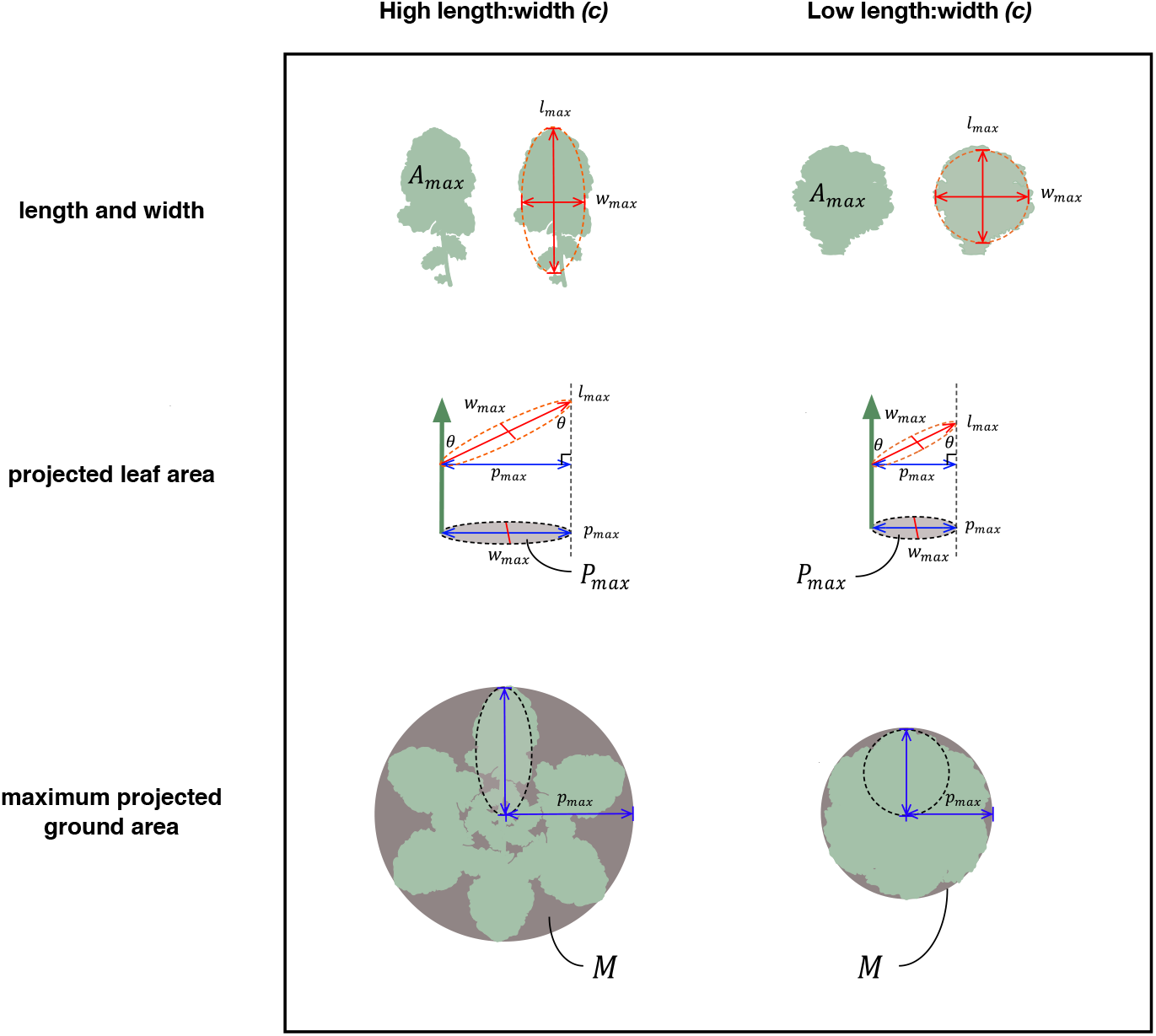
Concepts underlying leaf area index computation from single leaves. Shown here are measures used for calculating the denominator of leaf area index, maximum projected ground area *(M)*, based on projection of maximally sized leaves. Computation for the numerator are analogous, but uses dynamic values of *l, A,* and *p* rather their maximum. Differences in *M* may be achieved across genotypes that differ in the parameter, *c*, length-to-width ratio (unitless), given the same max leaf area, *A_max_*, and insertion angle, 0. Smaller *M* leads to greater rates of leaf area index increase.

Allocation to stem and root growth utilizes original TREES functions but is informed by new input parameters, root:shoot and leaf:stem ratios (**Table 1**), to reflect the diverse allocation strategies found across *B. rapa* morphotypes.

### Stochastic versus deterministic mode

The updated TREES model provides an option to simulate leaf growth in deterministic or stochastic mode via the input option *useLeafGamma.* If *useLeafGamma* is set to 0, relative rate of growth for every leaf that emerges during the simulation is informed by the same set of parameters. If *useLeafGamma* is set to 1, each leaf is informed by different parameter sets sampled from distributions informed by genotype-specific inputs, giving rise to stochastic growth.

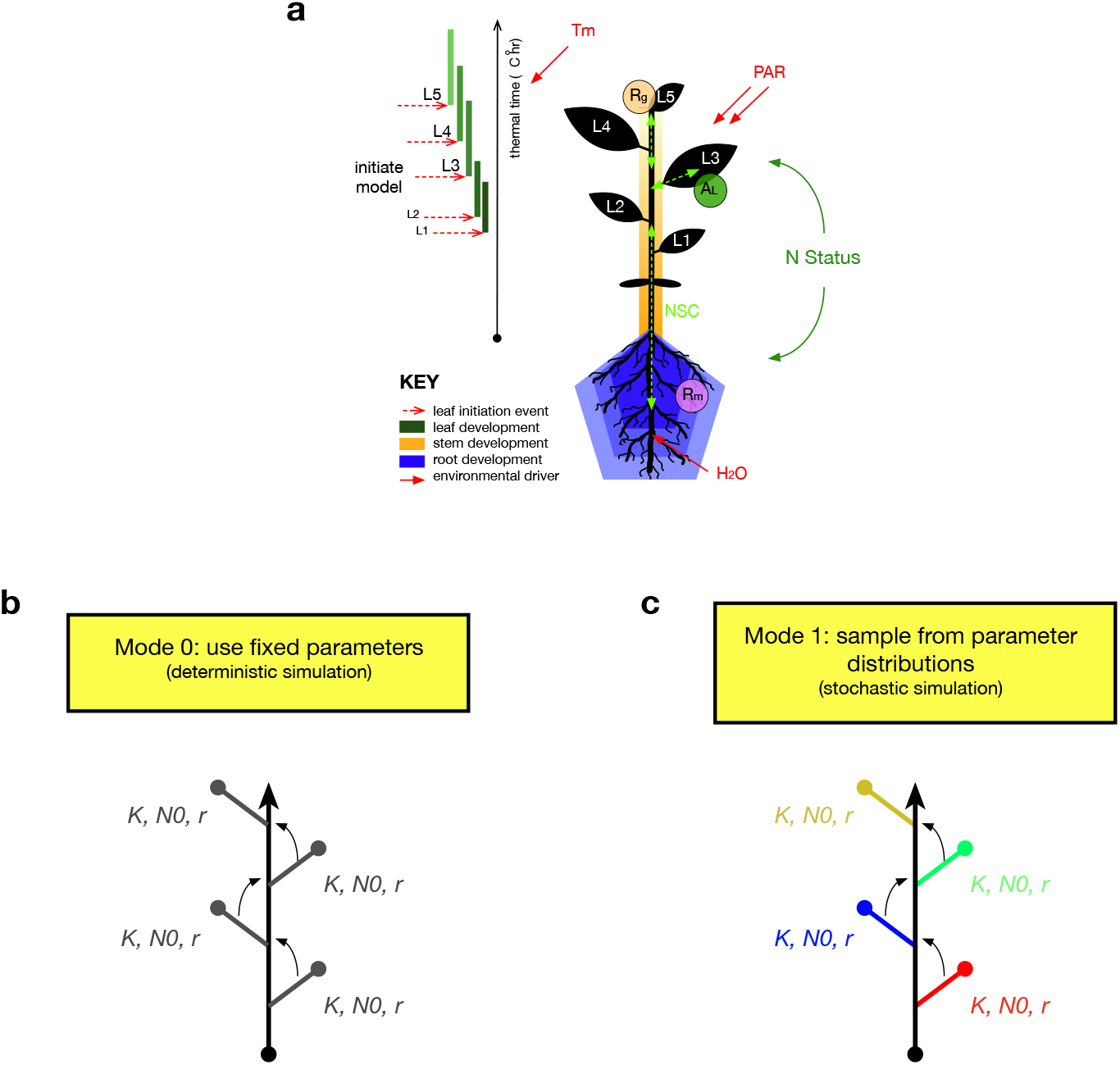
Modes for leaf growth for new TREES module. **(a)** Vegetative growth is modeled as sequential leaf development (green rectangles) through time from initiation of the first true leaf until transition to reproductive phase. Root and stem development (blue and orange shapes) are informed by progress of leaf development. Lighter shades for leaf, stem, and root development represent younger tissue. Panels **(b)** and **(c)** depict concepts underlying the two leaf growth modes in the new TREES leaf development module. In deterministic simulations, each leaf is parameterized by the same set of *K, N_0_*, and *r* values, represented by grey text in **(b)**. In stochastic simulations, each leaf is parameterized by a random sample out of *K, N_0_*, and *r* distributions parameterized by their rate and shape parameters. These random parameter sets are reflected in the different colors of *K, N_0_*, and *r* sets in **(c)**.

